# Companion cells with high florigen production express other small proteins and reveal a nitrogen-sensitive *FT* repressor

**DOI:** 10.1101/2024.08.17.608367

**Authors:** Hiroshi Takagi, Shogo Ito, Jae Sung Shim, Akane Kubota, Andrew K. Hempton, Nayoung Lee, Takamasa Suzuki, Jared S. Wong, Chansie Yang, Christine T. Nolan, Kerry L. Bubb, Cristina M. Alexandre, Daisuke Kurihara, Yoshikatsu Sato, Yasuomi Tada, Takatoshi Kiba, Jose L. Pruneda-Paz, Christine Queitsch, Josh T. Cuperus, Takato Imaizumi

## Abstract

The precise onset of flowering is crucial to ensure successful plant reproduction. The gene *FLOWERING LOCUS T* (*FT*) encodes florigen, a mobile signal produced in leaves that initiates flowering at the shoot apical meristem. In response to seasonal changes, *FT* is induced in phloem companion cells located in distal leaf regions. Thus far, a detailed molecular characterization of the *FT*-expressing cells has been lacking. Here, we used bulk nuclei RNA-seq and single nuclei RNA (snRNA)-seq to investigate gene expression in *FT*-expressing cells and other phloem companion cells. Our bulk nuclei RNA-seq demonstrated that *FT*-expressing cells in cotyledons and true leaves showed differences especially in *FT* repressor genes. Within the true leaves, our snRNA-seq analysis revealed that companion cells with high *FT* expression form a unique cluster in which many genes involved in ATP biosynthesis are highly upregulated. The cluster also expresses other genes encoding small proteins, including the flowering and stem growth inducer FPF1-LIKE PROTEIN 1 (FLP1) and the anti-florigen BROTHER OF FT AND TFL1 (BFT). In addition, we found that the promoters of *FT* and the genes co-expressed with *FT* in the cluster were enriched for the consensus binding motifs of NITRATE-INDUCIBLE GARP-TYPE TRANSCRIPTIONAL REPRESSOR 1 (NIGT1). Overexpression of the paralogous *NIGT1.2* and *NIGT1.4* repressed *FT* expression and significantly delayed flowering under nitrogen-rich conditions, consistent with NIGT1s acting as nitrogen-dependent *FT* repressors. Taken together, our results demonstrate that major *FT*-expressing cells show a distinct expression profile that suggests that these cells may produce multiple systemic signals to regulate plant growth and development.

## Introduction

Plants determine the seasonal timing of flowering based on environmental cues such as day length and temperature (1–6). The small protein FLOWERING LOCUS T (FT) is a florigen, a mobile signaling molecule that promotes flowering (7–10). In *Arabidopsis thaliana*, some phloem companion cells residing in the distal parts of leaves highly express *FT* (11, 12). Although *FT* exhibits only a single expression peak in the evening under common laboratory long-day (LD) conditions, *FT* is highly expressed in the morning and evening under natural light conditions (13, 14). The discrepancy between natural and laboratory conditions can be attributed to differences in the red-to-far-red light (R/FR) ratios. Adjusting the R/FR ratio to that observed in nature is sufficient to recreate the bimodal expression pattern of *FT* in the lab (13, 14).

Despite extensive research on the genetic regulation of *FT* expression and function, our understanding of the cells that express *FT* is limited (6). Although *FT* is expressed in some phloem companion cells, the precise molecular features that distinguish *FT*-expressing phloem companion cells from other companion cells have remained elusive (6). Recently, to understand the distinct features of *FT*-expressing cells, we employed Translating Ribosome Affinity Purification (TRAP)-seq, a method that identifies ribosome-associated transcripts in specific tissues or cell types (15). We found that the list of differentially translated transcripts in *FT*-expressing cells partially overlapped with that in the general phloem companion cell marker gene *SUC2*-expressing cells but also contained unique transcripts to *FT*-expressing cells (15). This supports the notion that the *FT*-expressing cells are similar but different from general phloem companion cells (11, 12). Our TRAP-seq datasets showed that several *FT* positive and negative transcriptional regulators and proteins involved in FT protein transport were specifically enriched in *FT*-expressing cells. We also found that a gene encoding the small protein FPF1-LIKE PROTEIN 1 (FLP1) is highly expressed in *FT*-expressing cells under LD conditions with adjusted R/FR ratio (hereafter, LD+FR conditions), but not under standard laboratory LD conditions (15). Further analysis revealed that FLP1 promotes flowering and initial inflorescence stem growth, suggesting that it orchestrates flowering and inflorescence stem growth during the transition from the vegetative to the reproductive stage.

Although our bulk tissue/cell-specific TRAP-seq analysis revealed some unique characteristics of *FT*-expressing cells, we still do not know their characteristics in a single-cell resolution. As *FT* is expressed at specific times of the day in relatively small numbers of phloem companion cells, we needed a method to collect the information from these rare *FT*-expressing cells in a time-dependent manner. Single-cell RNA-seq (scRNA-seq) is a powerful tool to study different cell populations within the complexity of biological tissues. In plant studies, especially for root tissues, the most popular procedure involves protoplasting of cells and the application of Drop-seq in the 10X genomics pipeline. However, protoplasting is not ideal for isolating highly embedded phloem companion cells efficiently (16). On top of this problem, our target is *FT*-expressing companion cells at the specific time point of the *FT* morning peak, thus the time required for tissue isolation must be short, so that the isolated cells still restore the time-of-day information. To overcome these technical challenges, we deployed single-nucleus RNA-seq (snRNA-seq) combined with fluorescence-activated nuclei sorting (FANS) to isolate GFP-labeled cell-type-specific nuclei at a specific time of the day. Here, we report the unique characteristics of *FT*-expressing cells at a single-cell level and identify new, growth condition-specific transcriptional repressors of *FT*.

## Results

### Bulk nuclei RNA-seq analysis finds profound expression differences in *FT*-expressing cells between cotyledons and true leaves

To investigate gene expression in the nuclei of specific cell populations, we used *Arabidopsis* transgenic lines expressing the gene encoding NTF [Nuclear Targeting Fusion protein consists of the nuclear envelope-targeting WPP domain, green fluorescent protein (GFP) and biotin ligase recognition peptide (BLRP) (17)] under the control of cell type-specific promoters **(Fig. S1A)**. In addition to the previously generated *pFT:NTF* line (15), we generated transgenic lines with *pSUC2* to capture most phloem companion cells, *pCAB2* for mesophyll cells, and *p35S* as a non-specific control (**Fig. 1A, Fig. S1A**). Although our original intention was the utilization of the INTACT (isolation of nuclei tagged in specific cell types), the method to capture tissue-specifically biotinylated nuclei using magnetic beads (17), we instead employed FANS to isolate the respective GFP-positive nuclei (18).

**Figure 1.**
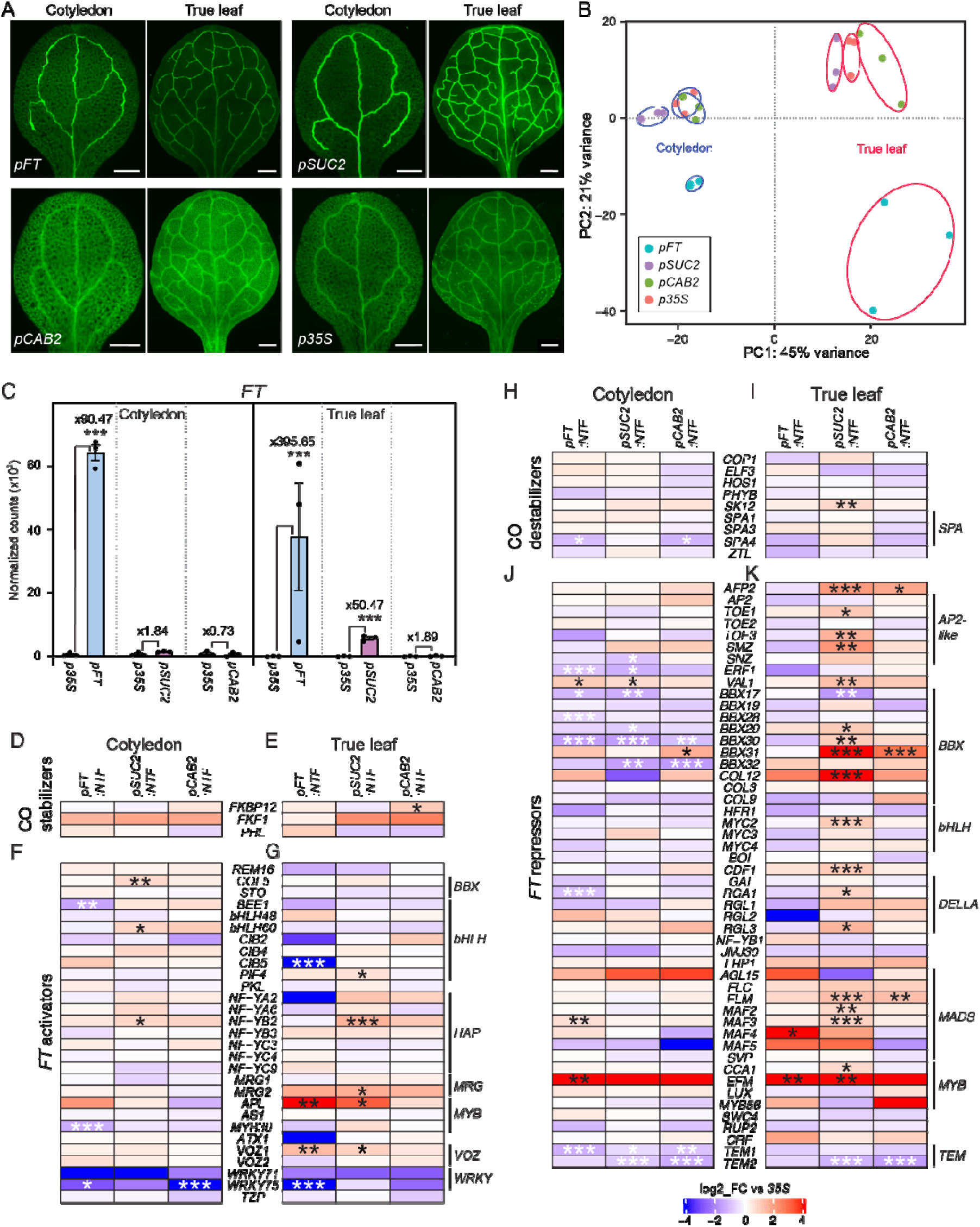
Tissue-and cell-type-specific gene expression in cotyledons and true leaves. (A) Representative images of ClearSee-treated cotyledons and true leaves of *pFT:NTF*, *pSUC2:NTF*, *pCAB2:NTF* and *p35:NTF* lines. Scale bar, 500 µm. (B) The first two principal components of bulk RNA-seq analysis for three independent cotyledon and true leaf samples. Cotyledon and true leaf samples are circled in blue and red, respectively. (C) DEseq2-normalized counts of *FT* transcripts in sorted nuclei for the *pFT:NFT*, *pSUC2:NFT,* and *pCAB2:NFT* lines compared to the *p35S:NTF* line. Fold enrichments of *FT* transcripts in each *NTF* line compared with those in *p35S:NTF* line are indicated. \*\*\**padj*<0.001. (D–K) Expression of genes encoding CO stabilizers (D and E), *FT* activators (F and G), CO destabilizers (H and I), and *FT* repressors (J and K) in cotyledons (D, F, H and J) and true leaves (E, G, I and K). Genes encoding proteins belonging to the same family were clustered in the heatmap. Bar color indicates log2-scaled fold-change relative to the *p35S:NTF* line. Asterisks denote significant differences from *p35S:NTF* (\**padj*<0.05; \*\**padj* <0.01; \*\*\**padj* <0.001). CO was removed from the analysis due to insufficient reads at ZT4.

Although *FT* is expressed in both cotyledons and true leaves, it was heretofore unknown whether *FT*-expressing cells in cotyledons and true leaves were equivalent to one another at the whole transcriptome level or showed tissue-specific differences in transcription. The previous histological analyses of the *Arabidopsis pFT:GUS* reporter line showed that *FT* expression in true leaves was confined to the distal part of the leaf vasculature, whereas in cotyledons, *FT* expression was observed in the broader part of the vein (11, 19), suggesting different profiles of *FT*-expressing cells in cotyledons and true leaves. To exclude possible tissue-specificity as a source of noise in our single-nucleus data, we conducted bulk nuclear RNA-seq of GFP-positive nuclei isolated from either cotyledons or true leaves. To do so, we harvested cotyledons and the first set of two true leaves from two-week-old transgenic plants grown under LD+FR conditions at Zeitgeber time 4 (ZT4), the time of the *FT* morning expression peak, and isolated GFP-positive nuclei within an hour using a cell sorter (**Fig. S1A, C**). We successfully collected GFP-positive nuclei from both cotyledons and true leaves for all transgenic lines (**Fig. S2**) and conducted bulk RNA-seq.

The principal component analysis (PCA) of the resulting expression data indicated separation by tissue and targeted cell population (**Fig. 1B**). Interestingly, the biggest difference among these datasets was the difference between cotyledons and true leaves, rather than a cell/tissue-type difference. Among different cells/tissues, *FT*-expressing cells show a distinct gene expression profile compared with other cell/tissue types examined.

Next, to ensure that we had properly enriched for the targeted cell populations, we conducted pairwise comparisons between the *p35S:NTF* control line and the cell type-specific *NTF* lines (**Data S1–6**). As expected, in true leaves of the *pSUC2:NTF* line, phloem companion cell marker genes *FT*, *SUC2*, *AHA3*, *APL*, *CM3,* and *AAP4* showed higher expression than in the *p35S:NTF* line, and the *pFT:NTF* line showed strong upregulation of *FT* in both cotyledon and true leaves (**Fig. 1C, Fig. S3A–E**). True leaves of the *pFT:NTF* line also showed upregulation of other phloem companion cell marker genes such as *AHA3*, *APL,* and *AAP4*; however, cotyledons of the *pFT:NTF* line only showed upregulation of *AAP4* (**Fig. S3A–E**), indicating some of known vascular marker genes are more suitable for representing vascular tissues in true leaves.

Similarly, in the *pSUC2:NTF* line, we observed differences in cell marker gene expression between cotyledons and true leaves. In cotyledons of the *pSUC2:NTF* line, *CM3* and *AAP4* were upregulated, but not *SUC2*, *FT*, *AHA3* and *APL*. To ensure that FANS properly enriched nuclei from *pSUC2:NTF* cotyledons, we checked the spatial expression pattern of each gene that was highly expressed in *pSUC2:NTF* cotyledons [adjusted *P*-value (*padj*) < 0.05, fold-change > 2-fold, **Table S1**]. The majority of the genes upregulated in the cotyledons of our *pSUC2:NTF* line were found to be expressed in the vasculature. As expected, expression of mesophyll marker gene *CAB2* was significantly lower in *pSUC2:NTF* cotyledon samples (*padj* = 1.5 x 10^-4^), in addition to the mesophyll cell marker *RBCS1A* in true leaf samples, suggesting that a significant amount of mesophyll cells was successfully removed by FANS (**Fig. S3F, G**).

Having ensured proper enrichments of the targeted nuclei, we compared global changes in expression among the tested transgenic lines. Consistent with the PCA analysis, the vast majority of differentially expressed genes within a given transgenic line did not overlap between cotyledons and true leaves (**Fig. S4, S5**). Next, we compared the expression of known *FT* positive and negative regulators in cotyledons and true leaves of the *pFT:NTF, pSUC2:NTF,* and *pCAB2:NTF* lines (**Fig. 1D–K**) (6).

Among the transcription factors that promote *FT* expression, *APL* and *VOZ1* were upregulated in true leaves of the *pFT:NTF* and *pSUC2:NTF* lines (**Fig. 1G**). In true leaves of the *pSUC2:NTF* line, a CO destabilizing factor was significantly upregulated (**Fig. 1I**), in addition to TFs that repress *FT* expression (**Fig. 1K**). In contrast to *pSUC2:NTF* true leaves, true leaves of the *pFT:NTF* line did not show significant upregulation for most of these *FT* repressors. This lack of *FT* repression appears to be crucial for specifying *FT*-expressing cells. In contrast, there were far fewer expression differences in cotyledons between the *pFT:NTF* and *pSUC2:NTF* lines, indicating that the respective cells in true leaves have diverged more than comparable ones in cotyledons.

Similarly, *FT* expression and FT transport appeared to be more tightly associated in true leaves than in cotyledons. As reported, the expression of the FT transporter gene *FTIP1* is highly associated with *FT* expression (15). Consistent with this earlier finding, among FT transporter genes, we observed significant upregulation of *FTIP1* expression in true leaves of the *pSUC2:NTF* and the *pFT:NTF* lines but not in the respective cotyledon samples (**Fig. S3H–K**).

To further support true leaves as most suitable for investigating *FT*-expressing cells, we performed Terms enrichment analysis with Metascape (20). In the *pFT:NTF* line, the genes involved in the “regulation of reproductive process” were upregulated in both cotyledons and true leaves, while the genes related to the term “long-day photoperiodism, flowering” were only upregulated in the true leaf samples of the *pFT:NTF* line (**Fig. S4**).

### snRNA-seq revealed subpopulation of phloem companion cells that highly express *FT*

Next, we used true leaves of the *pFT:NTF* and *pSUC2:NTF* lines for snRNA-seq using the 10X Genomics Chromium platform (**Fig. S1B**). We captured a total of 1,173 nuclei for the *pFT:NTF* line and 3,650 nuclei for the *pSUC2:NTF* line. In total, we captured 4,823 nuclei and detected 20,732 genes, a median of 149 genes per nuclei (**Fig. 2A, B**). Gene expression of these nuclei were projected in a two-dimensional UMAP as 11 different clusters (21) (**Fig. 2**). The nuclei in clusters 8 and 10 showed substantially higher number of transcripts than other clusters **(Fig. S6A and B**). Because this discrepancy made comparisons difficult, we focused on analyzing gene expression in the other clusters.

**Figure 2.**
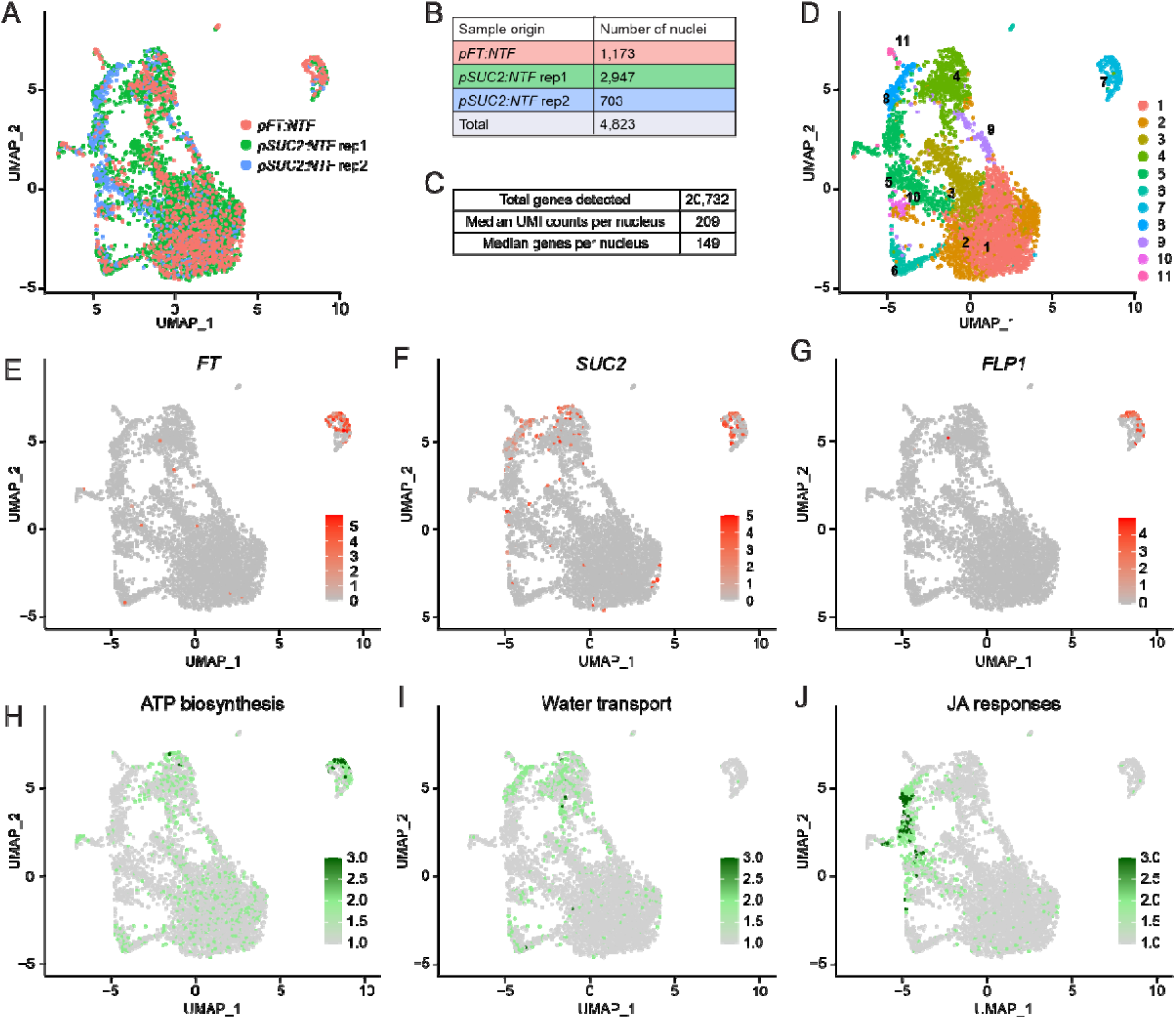
Single-nucleus RNA-seq identifies distinct subpopulations of phloem companion cells. (A) UMAP with the origin of nuclei indicated by color. (B and C) Table with the number of nuclei originated from each line and sample (B) and median UMI and number of genes detected per nucleus (C). (D) Seurat defined clusters with cluster numbers indicated. (E–G) UMAP annotated with normalized read counts for *FT* (E), *SUC2* (F), and *FLP1* (G) expression. (H–I) UMAP annotated with average read counts of genes related to ATP biosynthesis (H), water transport (I), and JA responses (J). Color bars indicate gene expression levels. For the lists of genes, see Fig. S7D (pink highlighted), S8E and S9B.

We first asked which nuclei highly expressed *FT*. These nuclei resided in cluster 7 (**Fig. 2D**, **2E**). *SUC2* expression showed a far broader distribution across clusters than *FT* expression (**Fig. 2F**). The nuclei in cluster 7 significantly highly expressed 268 genes compared with total population (**Data S7**). Among these genes, we found *FLP1* (**Fig. 2G**), a gene encoding a flowering-promoting factor acting in parallel with FT (15). Next, we cross-checked these 268 genes with genes identified in the previous TRAP-seq experiment (15). 202 of the 268 genes showed higher expression in *FT*-expressing cells (*pFT:FLAG-GFP-RPL18*) than in whole companion cells (*pSUC2:FLAG-GFP-RPL18*) (**Fig. S7A**) in the prior study, validating our single nuclei analysis.

Further, *FKBP12*, a gene encoding a CO stabilizing protein was expressed in cluster 7 (**Fig. S7B**).

To explore the *FT*-expressing cluster 7, we conducted Terms enrichment analysis using Metascape (20). Genes involved in “Oxidative phosphorylation” were enriched in cluster 7 (**Fig. S7C**), suggesting that ATP synthesis is particularly active in nuclei belonging to this cluster (**Fig. 2H**). Genes involved in proton-generating complex I–IV and ATP synthesizing complex V in mitochondrion membrane were also upregulated in cluster 7 (**Fig. S8D**), suggesting that the entire ATP synthesis pathway is activated in *FT*-expressing nuclei. Additional terms such as “generation of precursor metabolite and energy” and “purine nucleotide triphosphate metabolic process,” further indicated upregulation of ATP production. Since phloem companion cells actively hydrolyze ATP to generate proton gradients to load sucrose and amino acids (22), it is plausible that *FT*-expressing phloem companion cells generate a substantial amount of ATP for transport.

We next inquired about the characteristics of other phloem nuclei clusters. Like cluster 7, cluster 4 showed significantly higher *SUC2* expression compared with other clusters, suggesting companion cell identity (**Fig. 2F**, **Data S7**). Moreover, the genes *LTP1*, *MLP28*, and *XTH4* were upregulated in cluster 4, consistent with prior studies showing their exclusive expression in vascular tissues (particularly in the main vein) that include phloem (**Fig. S8A–C**) (23–25). Terms enrichment analysis found that genes encoding aquaporin were enriched in cluster 4 (**Fig. 2I, Fig. S8D, E**). To understand which part of leaf tissue consists of cluster 4 cells, we marked the promoter activity of *PLASMA MEMBRANE INTRINSIC PROTEIN 2.6* (*PIP2;6*), an aquaporin gene highly expressed in cluster 4, using the GFP-fusion protein **(Fig. S8F)**. It showed the strongest signal in the main vein, consistent with the previous GUS reporter assay (26). Taken together, the nuclei in cluster 4 are likely to be derived from phloem cells in the main vein that are active in solute transport.

Cluster 5 is mostly composed of nuclei from the *pSUC2:NTF* line (464 out of 515 nuclei) (**Fig. S6B**). The nuclei in this cluster showed differentially expressed genes related to defense (**Fig. S9A**). Specifically, *MYC2,* a master regulator of response to jasmonic acid (JA), and the JA biosynthetic genes *LOX3*, *LOX4*, *OPR3,* and *OPCL1,* are expressed in cluster 5 (**Fig. S9B**). These genes are known to be expressed in vascular tissues, with *OPR3* expressed in phloem companion cells and *LOX3*/*4* in phloem (27–29). This subpopulation of phloem cells appears to play pivotal roles in JA-dependent defense responses. We did not include a nuclei fixation step in our FANS protocol, because we encountered tissue/nucleus clumping with fixation, which caused an issue in the sorting process. Because these cells were not fixed, the nuclei isolation process may have triggered wounding responses due to chopping samples before the sorting process (**Fig. S1B**).

Cluster 6 consisted of both phloem parenchyma and bundle sheath cells (**Fig. S10A–F**). Subclustering revealed that the nuclei in the upper part of cluster 6 expressed phloem parenchyma marker genes, and the nuclei in the lower part expressed bundle sheath marker genes (16) (**Fig. S10C–F**). Overall, the genes expressed in this cluster were enriched for genes functioning in sulfur metabolisms (**Fig. S10G**), including glucosinolate biosynthesis (**Fig. S10H**). A previous study showed that sulfur metabolic and glucosinolate biosynthetic genes are actively expressed in bundle sheath cells (30).

Lastly, Clusters 1, 2, and 3 showed expression of mesophyll cell marker genes **(Fig. S6C–F)**, suggesting that some mesophyll cells were included during the sorting process. This result might be due to weak induction of the *FT* and *SUC2* promoters in mesophyll cells (15). Indeed, we found both genes to be expressed at low levels in mesophyll protoplasts **(Fig. S11)**. In summary, using snRNA-seq combined with FANS, we revealed the presence of a unique subpopulation of cells with high nuclear *FT* expression. In these nuclei, genes related to ATP synthesis were enriched. Moreover, we identified other phloem cell clusters enriched for genes in water transport and JA response, which were not previously reported (16).

Next, we subclustered the nuclei in cluster 7, finding three subclusters (**Fig. 3**). Subcluster 7.2 contained the nuclei with the highest *FT* expression, in addition to expression of the companion cell marker genes, *SUC2* and *AHA3*. These nuclei contained fewer transcripts of the mesophyll marker genes *RBCS1A*, *CAB3,* and *CAB2* (**Fig. 3C, D**), consistent with a prior scRNA-seq study showing the negative correction between the expression of vasculature and mesophyll cell marker genes (31). The vascular tissue located at the distal part of true leaves, where *FT* is expressed highly, is developmentally old, whereas veins emerging from the bottom part of leaves are developmentally young (32). Thus, subcluster 7.3 may be comprised of nuclei from developmentally younger companion cells that have not fully matured yet, whereas those in subcluster 7.2 are older and have gained a stronger companion cell identity.

**Figure 3.**
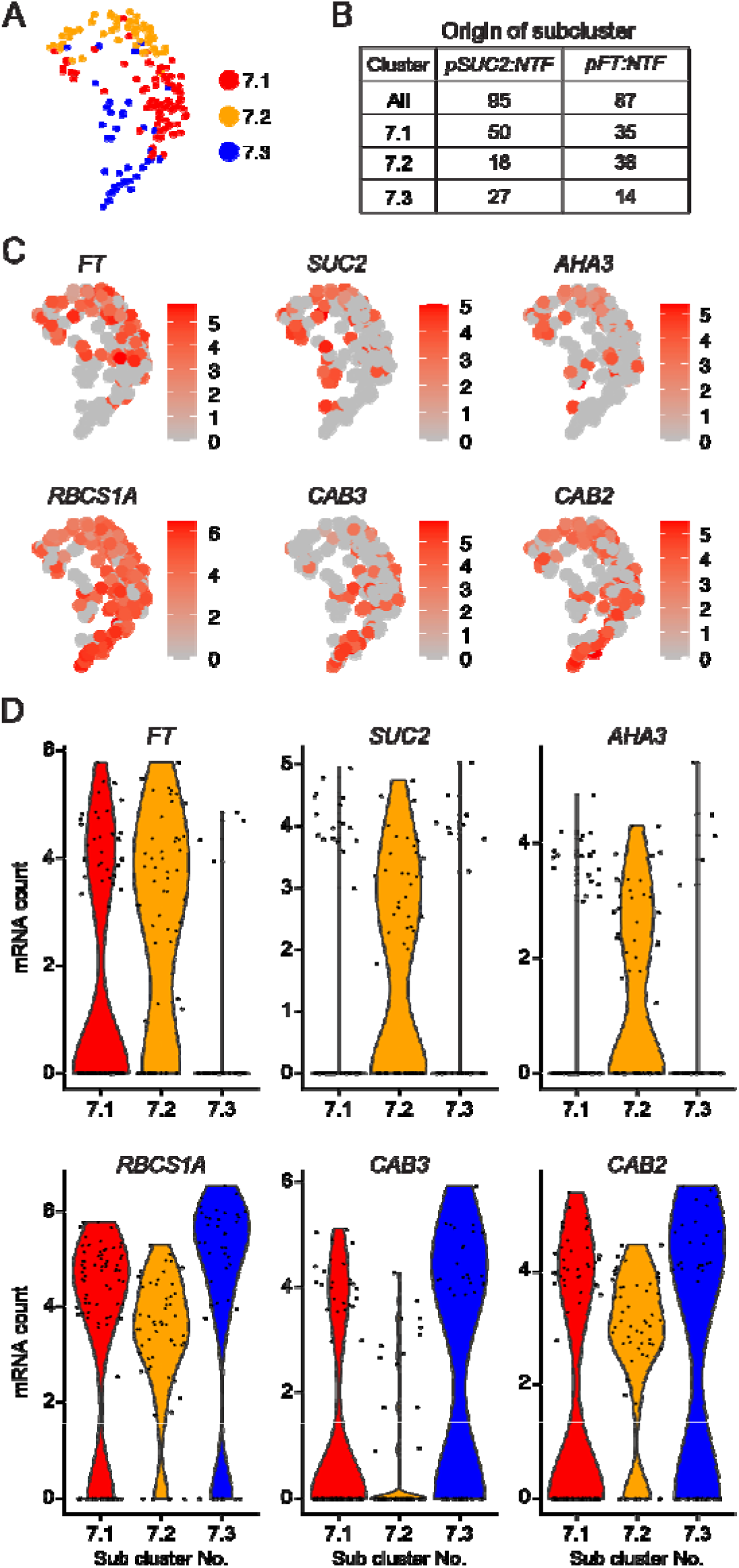
Phloem companion cell and mesophyll cell marker gene expression in cluster 7. (A) Subclustering of cluster 7. Colors indicate each subcluster. (B) Table showing the number of nuclei originated from each *NTF* line in cluster 7. (C) UMAP of normalized read counts of companion cell marker genes (*FT*, *SUC2*, and *AHA3*) and mesophyll cell marker genes (*RBCS1A*, *CAB3*, and *CAB2*). (D) Violin plot of normalized read counts of marker genes for companion cells and mesophyll cells across the three subclusters.

### Phloem companion cells with high *FT* expression express other genes encoding small proteins

In LD+FR conditions, *FT*-expressing cells also express *FLP1,* which encodes another small protein with systemic effects on flowering and inflorescent stem growth (15). We asked whether the cluster 7 nuclei might express other genes encoding small proteins, which could move through phloem flow. Indeed, the median number of amino acids encoded in the differentially expressed genes of cluster 7 is smaller than those in clusters 4 and 5 **(Fig. 4A)**. To validate whether high *FT*-expressing vascular cells co-express genes specifically upregulated in cluster 7, which encode small proteins, we selected eight such genes and generated their respective promoter fusions with *H2B-tdTomato* in the *pFT:NTF* background. The spatial expression patterns of these eight genes and high *FT* signals overlapped (**Fig. 4B, S12**). *FT* and cluster 7-enriched genes examined tended to express highly in the distal part of minor veins, but weakly in the main vein adjacent to the petiole, showing a clear difference from the cluster 4-enriched gene *PIP2;6* that is expressed highly in the main vein (**Fig. S8F, S12**). It should also be noted that the spatial expression patterns of *FT* and other cluster 7-enriched genes did not completely overlap; there were cells expressing only one of them at high levels (**Fig. S12, S13**).

**Figure 4.**
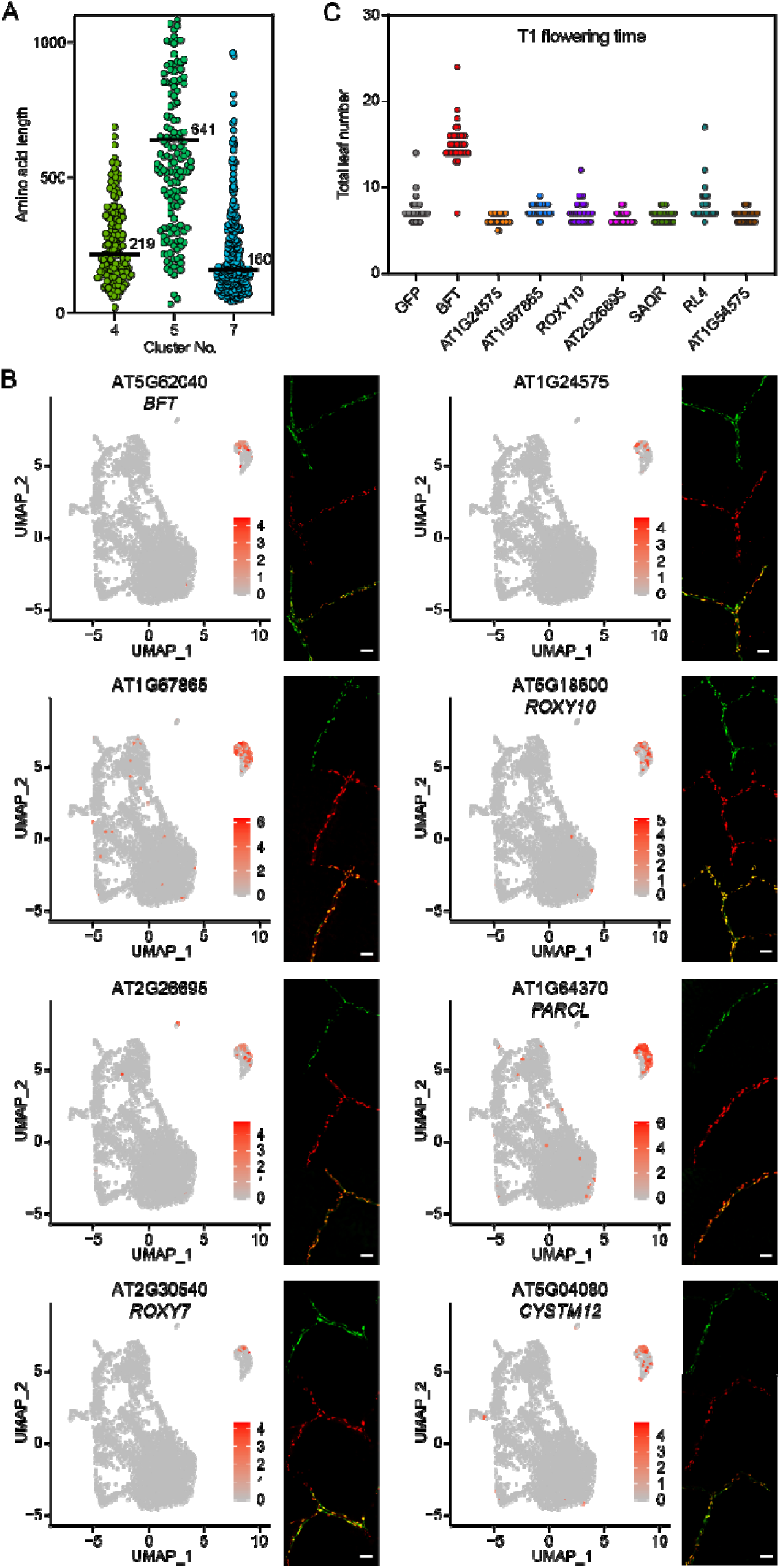
*FT*-expressing cells express genes encoding other small proteins. (A) Amino acid length of proteins encoded by genes differentially expressed in clusters 4, 5, and 7. Black bars indicate the median amino acid length. (B) Expression of the *pFT:NFT* line (green) and promoter fusions of selected cluster 7 genes with *H2B-tdTomato* (red) in true leaves. The selected genes were differentially expressed in cluster 7 and encoded small proteins. Yellow color shows an overlap between green and red signals. Scale bar, 50 µm. (C) Flowering time measurements of T1 transgenic plants overexpressing selected cluster 7 genes driven by the *pSUC2* promoter. Eight genes were tested, five of which were tested for overlap with *FT* expression in (B).

To assess the biological significance of *FT* generated from cluster 7, we knocked down *FT* transcripts using an artificial microRNA (amiRNA) targeted to *FT* (*amiR-ft*) controlled by the promoter of cluster 7-enriched *ROXY10* gene. As a comparison, we also expressed the *amiR-ft* gene using *SUC2* (general companion cell-marker gene), *PIP2;6* (cluster 4-enriched gene), and *GC1* [a marker gene for guard cells where *FT* is also expressed (33)], and tested the effect on the timing of flowering at the T_1_ generation. If *FT* generated from cluster 7 is important, as *ROXY10* is specifically expressed in cluster 7, late flowering of *pROXY10:amiRNA-ft* is anticipated. As a result, *pROXY10:amiRNA-ft* and *pSUC2:amiR-ft* lines severely delayed flowering (in fact, *pSUC2:amiRNA-ft* flowered later than *ft-101* mutant likely due to the effect of the hygromycin selection plate). On the other hand, *pPIP2;6:amiR-ft* was comparable with the negative control *pGC1:amiRNA-ft* line **(Fig. S14A)**. The basal expression level of *ROXY10* was much lower than that of *SUC2* and *PIP2;6* **(Fig. S14B)**; therefore, the delayed flowering phenotype of *pROXY10:amiR-ft* lines was not attributed to the promoter strength of genes used to drive the amiRNA. These results indicate that *FT* expressed in a relatively small but unique cluster 7 is biologically significant for flowering time regulation, although *FT* expressed in other companion cells other than cluster 7 cells may also contribute to flowering.

Next, we asked whether some of the genes enriched in cluster 7 encode new systemic flowering or growth regulators. We selected eight genes, five of which were tested for overlap with *FT* expression (**Fig. 4B**) and overexpressed each one of them using the *SUC2* promoter. Some of these genes were previously implicated in flowering. *BFT* is an *FT* homolog and acts as a floral repressor. *SENESCENCE-ASSOCIATED AND QQS-RELATED* (*SAQR*) promotes flowering under short-day (SD) conditions (34). *RAD-LIKE 4* (*RL4*) has high homology to the *RADIALIS* (*RAD*) gene in *Antirrhinum majus*, whose ectopic expression in *Arabidopsis* causes growth defects and late flowering (35). We measured the flowering time of the overexpression lines in independent T_1_ plants (**Fig. 4C**). T_1_ *pSUC2:BFT* plants grown under LD+FR conditions showed severely delayed flowering compared with *pSUC2:GFP* control lines (**Fig. 4C**). This result is in stark contrast to prior results obtained under standard laboratory light conditions (36), suggesting that BFT proteins might be mobile or more functionally active under natural light conditions. None of the other transgenic lines showed noticeable changes in flowering time.

### Motif enrichment analysis and transgenic studies identify *NIGT1* transcription factors as direct *FT* repressors

We performed motif enrichment analysis on the promoters of the 268 genes differentially expressed in cluster 7 (37, 38), using randomly selected 3,000 genes as a control set. The most enriched motif was the motif of the NITRATE-INDUCIBLE GARP-TYPE TRANSCRIPTIONAL REPRESSOR 1.2 (also known as HHO2, *P* = 9.3 x 10^-6^) (**Fig. 5A**; **Data S8**). Although the NIGT1/HHO transcription factors (TFs) are primarily known as repressors of genes involved in nitrate uptake, constitutive expression of *NIGT1.2* significantly decreases *FT* expression (39, 40). In fact, there are two potential NIGT1 binding sites located within and adjacent to the *shadow 1* and *2* domains (*S1 and S2*) in the *FT* regulatory region (41) (**Fig. S15A**). These domains are highly conserved among *Brassicaceae* species and important for *FT* transcriptional regulation.

**Figure 5.**
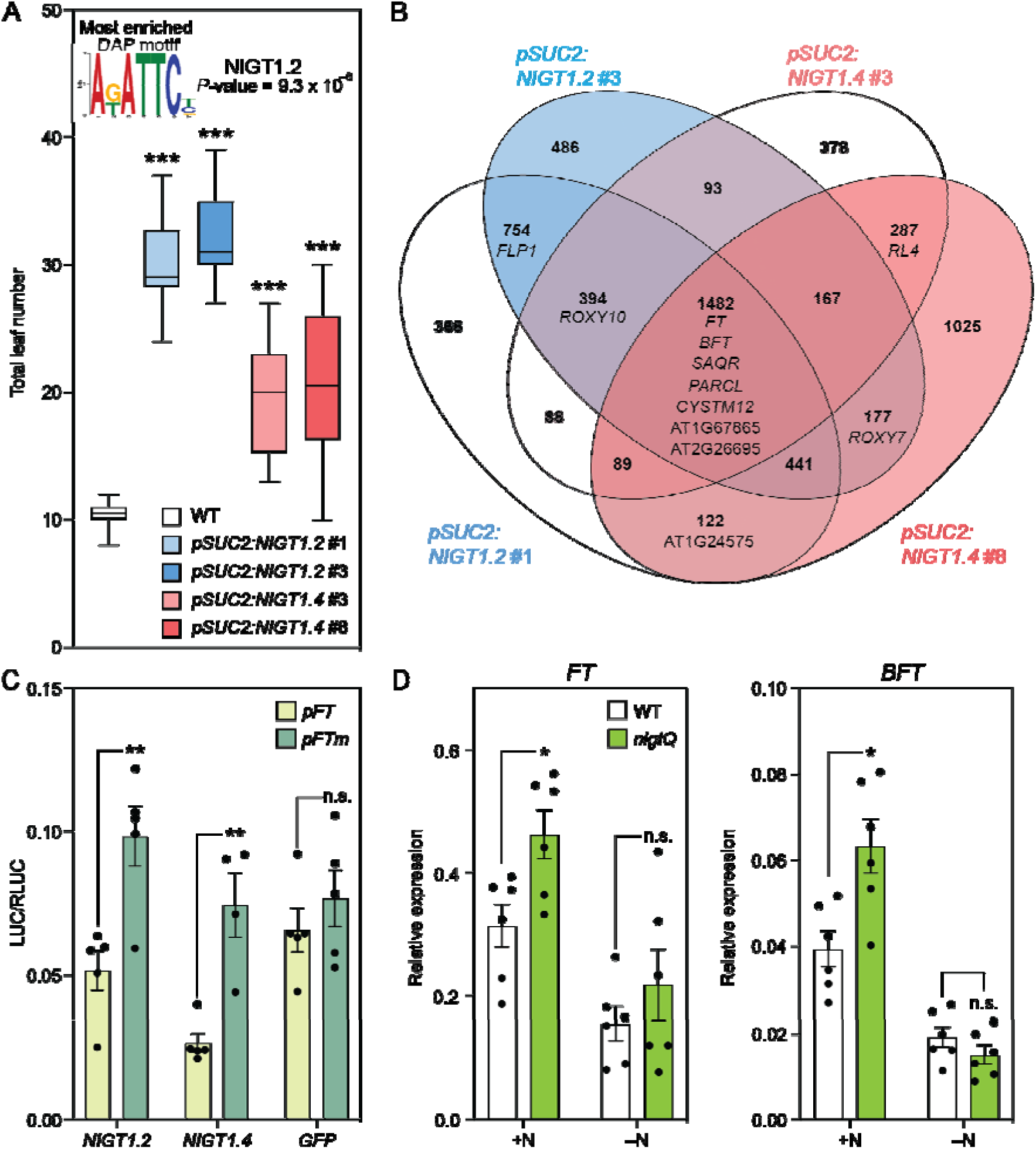
NIGT1 transcription factors are repressors of *FT*. (A) Motif enrichment analysis using the 268 genes differentially expressed in cluster 7 and flowering time of the *pSUC2:NIGT1.2* and *pSUC2:NIGT1.4* lines. The NIGT1.2 binding site was the most enriched motif in cluster 7 differentially expressed genes. The bottom and top lines of the box indicate the first and third quantiles, and the bottom and top lines of the whisker denote minimum and maximum values. The bar inside the box is the median value (*n* =16). (B) A Venn diagram showing the significantly down-regulated genes from bulk RNA-seq of two independent *pSUC2:NIGT1.2* and *pSUC2:NIGT1.4* lines. (C) The effect of *NIGT1.2*, *NIGT1.4* and *GFP* on wild-type and mutated *FT* promoter activity in tobacco transient assay. The 1 kb of *FT* promoter (*pFT*) and the same *FT* promoter with all NIGT1-binding sites mutated (*pFTm*) control the expression of firefly luciferase (*LUC*) gene. The LUC activity was normalized with Renilla luciferase (RLUC) activity controlled by *35S* promoter. The results are means ± SEM with each dot representing biological replicates (*n* = 4 or 5). (D) Relative gene expression levels of 14-day old wild-type (WT) and *nigtQ* seedlings at ZT4. Plants were grown with high nitrogen (+N) and without (–N). The results are means ± SEM with each dot representing biological replicates (*n* = 6). Asterisks denote significant differences from WT (\**P*<0.05, \*\**P*<0.01, \*\*\**P*<0.001, *t*-test). n.s. indicates not significant.

Moreover, the NIGT1 binding motifs reside in close proximity to the CO binding motif (TGTGNNATG, CO-responsive element, CORE) (41, 42), suggesting that NIGT1 binding could affect CO binding.

We performed a yeast one-hybrid (Y1H) screen of 1,957 TFs against four tandem repeats of a short *FT* promoter sequence containing the *S1* and *S2* domains and the two NIGT1 binding sites (43, 44). We found that NIGT1.1/HHO3, NIGT1.2/HHO2, NIGT1.3/HHO1, NIGT1.4/HRS1, and HHO5 specifically bind to this sequence within the *FT* regulatory region (**Data S9**). Next, we overexpressed five different *NIGT1/HHO* genes using the *SUC2* promoter. At T_1_ generation, we observed the late flowering phenotype in the *pSUC2:NIGT1.2* and *NIGT1.4* lines, while it appeared less clear in the *pSUC2:NIGT1.1*, *NIGT1.3*, and *HHO5* (data not shown). Therefore, we generated T_3_ lines of *pSUC2:NIGT1.2* and *pSUC2:NIGT1.4* and measured their flowering time.

Under LD+FR conditions, these transgenic plants flowered significantly later than wild-type plants (**Fig. 5A**). To test which genes were altered by the overexpression of *NIGT1.2* and *NIGT1.4*, we conducted RNA-seq analysis of the respective transgenic plants **(Fig. 5B, Data. S10–13)**. We found that genes specifically expressed in cluster 7 such as *FT*, *BFT*, and *SAQR* were downregulated in both the *pSUC2:NIGT1.2* and *pSUC2:NIGT1.4* lines. *FLP1* expression was decreased only in *pSUC2:NIGT1.2* lines, which might explain why *pSUC2:NIGT1.2* plants exhibited a more severe late flowering phenotype than *pSUC2:NIGT1.4* plants. Similar results were obtained with qRT-PCR analysis (**Fig. S16**). To test if NIGT1.2 and NIGT1.4 directly regulate the *FT* promoter activity, we conducted the tobacco transient promoter luciferase (LUC) assay using the *FT* promoter (*pFT*) and the one with the mutations in three potential NIGT1 binding sites (*pFTm*) **(Fig. S15B)**. Our data showed that co-expression of NIGT1.2 and NIGT1.4 significantly decreased the promoter activity of *pFT* compared with that of *pFTm*, while GFP did not change the *pFT* activity compared with *pFTm* **(Fig. 5C)**, suggesting that NIGT1s directly repress *FT* expression level. Consistent with our snRNA-seq data (**Fig. S17A, B**), previous studies reported that *NIGT1.2* is expressed broadly in shoots while *NIGT1.4* is expressed only in roots (39), suggesting that *NIGT1.2* is the major player in *FT* regulation. Interestingly, neither *NIGT1.2* or *NIGT1.4* expressed highly in cluster 7. We speculate that they have a role to prevent the misexpression of *FT* and perhaps other cluster 7 genes in non-cluster 7 cells.

Ample nitrogen prolongs the vegetative growth stage and delays flowering (45); however, the detailed mechanisms of this delay remain largely unknown (45). Since *NIGT1* genes are induced under high nitrogen conditions, we tested the quadruple mutant of the *NIGT1* genes (*nigtQ* mutant) for *FT* and *BFT* expression with (+N) and without high nitrogen (–N) (39). The *nigtQ* plants showed enhanced *FT* and *BFT* expression compared to the wild-type plants under +N but not –N conditions, suggesting that the NIGT1 TFs act as novel nitrate-dependent regulators of flowering (**Fig. 5D, S17C**). To test if *nigtQ* mutations affect flowering timing under different nitrogen conditions, we transplanted 9-day-old WT and *nigtQ* seedlings grown on Murashige and Skoog-based media with 20 mM NH_4_NO_3_ to new media containing either 20 mM or 2 mM NH_4_NO_3_. The *nigtQ* plants bolted earlier in days than WT when they grew with 20 mM NH_4_NO_3_, whereas the flowering time of WT and *nigtQ* was almost identical under 2 mM, suggesting that *NIGT1* genes delay flowering under ample nitrogen conditions (**Fig. S17D and E**). However, there was no difference between WT and *nigtQ* in leaf number at bolting (**Fig. S17F**). Therefore, it appears that the *nigtQ* mutation enhanced overall plant growth speed rather than developmental timing of flowering. We also have counted leaf numbers of the *nigtQ* at bolting on nitrogen-rich soil. The mutant generated slightly more leaves than WT when they flowered **(Fig. S17G)**. These results suggest that the NIGT-derived fine-tuning of *FT* regulation is conditional on higher nitrogen conditions.

## Discussion

Plants utilize seasonal information to determine the onset of flowering. One of the key mechanisms of seasonal flowering is the transcriptional regulation of the florigen *FT* in leaves. Our previous study revealed that *FT*-expressing cells also produce FLP1, another small protein that may systemically promote both flowering and elongation of leaves, hypocotyls, and inflorescence stems (15). Therefore, a more detailed characterization of *FT*-expressing cells might identify additional components that are essential for the integration of the many environmental cues in natural environments into the precise timing of flowering and related developmental changes. Here, we applied bulk nuclei and snRNA-seq and transgenic analysis to examine *FT*-expressing cells at high resolution.

### Differences in the spatial expression patterns of *FT* between cotyledons and true leaves are likely driven by negative regulators

In young *Arabidopsis* plants grown in LD, true leaves show *FT* expression in the distal and marginal parts of the leaf vasculature, while cotyledons express *FT* across entire veins (11, 19). We used bulk RNA-seq to examine the *FT*-expressing cells in cotyledons versus those in true leaves, using FANS-enriched nuclei. Tissue-specific differences (cotyledons vs. true leaves) in gene expression were more significant than those observed for the enriched cell populations, an unexpected result as cotyledons and true leaves are not typically treated as separate organs.

The true leaves but not the cotyledons of the *pSUC2:NTF* and *pFT:NTF* lines showed differences in the expression of *FT* negative regulators. This result suggests that *FT* expression is strongly repressed in most of the true leaf companion cells, contributing to the spatial expression pattern of *FT* observed in true leaves. In this study, we focused on the *FT* expression peak in the morning (ZT4) under LD+FR conditions; however, in the future, it would be interesting to investigate the expression patterns of *FT* regulators at the *FT* evening peak (ZT16), as our results suggest that there are two different mechanisms regulating each *FT* peak in natural long days (13, 14).

### A subpopulation of phloem companion cells highly expresses *FT* and other genes encoding small proteins

Our snRNA-seq identified a cluster with nuclei derived from cells with high *FT* expression that controls flowering time. The cluster 7 nuclei are derived from cells that appear to be metabolically active and produce ATP, which may be used to upload sugars, solutes, and small proteins, including FT, into the phloem sieve elements. Based on subclustering analysis, *FT* expression levels were positively correlated with companion cell marker gene expression and negatively correlated with mesophyll cell marker gene expression. The observed shift from mesophyll to companion cell identity may represent the trajectory of companion cell development (31).

In clusters 4 and 5, we found nuclei isolated from phloem cells with high expression of aquaporin and JA-related genes, respectively. These cell populations were not identified in a previous protoplast-based scRNA-seq analysis targeting phloem (16). We speculate that these cells are highly embedded in the tissue and therefore practically difficult to isolate. Previous studies show that JA biosynthetic genes such as *LOX3/4* and *OPR3* are highly expressed in phloem including companion cells (27, 29), consistent with our results.

Aquaporin genes such as *PIP2;1* and *PIP2;6* are thought to be highly expressed in leaf xylem parenchyma and bundle sheath tissues (26). However, cluster 6, which contains nuclei isolated from bundle sheath and phloem parenchyma cells, showed fewer expressed aquaporin genes than the cluster 4 nuclei, which also expressed *SUC2*. This result might indicate that the cells active in water transport exist not only in bundle sheath or xylem parenchyma cells but also in a subpopulation of phloem cells. In summary, *SUC2*/*FT*-expressing vasculature cells consist of cell subpopulations with different functions within true leaves.

A drawback of our snRNA-seq analysis was shallow reads per nucleus. It appears mainly due to the low abundance of mRNA in FANS-isolated nuclei from 2-week-old leaves. Based on our calculation, the average mRNA level per nucleus in our isolated samples is approximately 0.2 pg (3,000 pg mRNA from 15,000 sorted nuclei). Future technological advance is needed to improve the data quality.

### BFT may fine-tune the balance between flowering and growth as a systemic anti-florigen

The *FT*-expressing nuclei of cluster 7 preferentially expressed genes encoding small proteins including *FLP1*, *BFT,* and *SAQR* (34, 36, 46). Although a previous study reported that *BFT* driven by the *SUC2* promoter does not alter flowering time (36), our result shows that overexpression of this gene delays flowering under LD+FR conditions (**Fig. 4C**). This discrepancy is likely due to our use of growth conditions with the adjusted R/FR ratio that mimics natural light conditions, as the majority of our transgenic lines showed a similar late flowering phenotype.

Why do *Arabidopsis* plants produce florigen FT and anti-florigen BFT in the same cells? There is precedent for the co-expression of florigen and anti-florigen in leaves in other plants. In Chrysanthemums, a weak florigen *CsFTL1* and a strong anti-florigen *CsAFT* are both expressed in leaves in long days. Short-day conditions repress *CsAFT* but induce the expression of a strong florigen *CsFTL3* to initiate flowering (47, 48). The balance in florigen and anti-florigen expression is also important for wild tomatoes to control flowering time, and this regulation was lost in domesticated tomatoes, resulting in day-neutral flowering behaviors (49). In *Arabidopsis*, the FT-homolog floral repressor TERMINAL FLOWER 1 (TFL1) is expressed at the shoot apical meristem and competes with FT for physical interaction with FD (50). In addition to preventing precocious flowering, the presence of TFL1 is crucial for the balance between flowering initiation and stem growth because FT terminates inflorescence stem growth (51). Like FT and TFL1, BFT directly binds to FD (36), suggesting that BFT’s mode of action is similar to that of TFL1. However, unlike for TFL1, loss of BFT does not affect flowering time under standard growth conditions (36, 46). The lack of a visible flowering phenotype in the *bft* knockout mutant might be due to the substantially lower expression of *BFT* compared to *FT* under standard growth conditions (36). Consistent with this interpretation, the *bft* knockout mutant is significantly delayed in flowering under high salinity conditions where *BFT* gene expression is strongly induced (36).

Thus, BFT appears to act as an anti-florigen under specific growth conditions, possibly including LD+FR conditions. Given the complexity of natural environments, the existence of multiple specialized anti-florigens might facilitate signal integration and reduce noise in the onset of flowering and stem growth.

### NIGT1 TFs contribute to the nutrient-dependency of flowering time

To identify potential novel transcriptional regulators of *FT*, we identified enriched motifs in the promoters of the 268 genes that were differentially expressed in cluster 7. This analysis indicated that NIGT1 TFs may affect the expression of *FT* and other genes co-expressed with *FT*. Using Y1H screening, we found that all NIGT1s and HHO5 can bind to the enhancer sequences derived from the *FT* promoter. NIGT1s are involved in nitrogen absorption as negative regulators (39, 40). It is well-known that nitrogen availability affects the onset of flowering; however, most of the mechanistic underpinnings remain unknown. A recent study showed that ample nitrogen availability causes phosphorylation of FLOWERING BHLH 4 (FBH4), a direct positive regulator of CO, which results in attenuation of FBH4 transcriptional activity (45). This mechanism acts upstream of *FT*, whereas the NIGT1 TFs likely act as direct repressors of *FT*. Multiple nitrogen-dependent mechanisms acting on *FT* regulation might ensure the proper balance between nutrient availability and resource allocation toward developmental transitions.

## Materials and Methods

### Molecular cloning and plant materials

All *Arabidopsis thaliana* transgenic plants and mutants are Col-0 backgrounds. *ACT2:BirA* plant (17) was used as the *Arabidopsis* genetic background to generate *NTF* expressing lines. The *NTF* cDNA was controlled by the *35S* promoter or the tissue-specific promoters: *CAB2* (AT1G29920), *SUC2* (AT1G22710), and *FT* (AT1G65480). The *NTF* sequences were amplified using primers (5’-CACCATGGATCATTCAGCGAAAACCACACAG-3’ and 5’-TCAAGATCCACCAGTATCCTCATGC-3’) and cloned into the pENTR/D-TOPO vector (Invitrogen). The *CAB2* promoter region (324 bp) was amplified by the primers (5’-CACCATATTAATGTTTCGATCATCAGAATC-3’ and 5’-TTCGATAGTGTTGGATTATATAGGG-3’) and cloned into the pENTR 5’-TOPO (Invitrogen) (52). The *SUC2* promoter region (2,302 bp) was amplified by primers (5’-GGTGCATAATGATGGAACAAAGCAC-3’ and 5’-ATTTGACAAACCAAGAAAGTAAG-3’) and cloned into the pENTR 5’-TOPO. The *CAB2* and *SUC2* promoter sequences were fused with *NTF* in the binary GATEWAY vector R4pGWB501 through LR clonase reaction (53). The *CAB2:NTF* and *SUC2:NTF* constructs were transformed into wild-type Col-0. The *pFT:NTF* line was described in the previous study (15).

The *H2B-tdTomato* constructs containing promoter regions of cluster 7 highly expressed genes were made by swapping the heat-shock promoter (*pHS*) of *pPZP211 HS:H2B-tdTomato* (*15*). Promoter sequences were amplified by the forward primer containing SbfI site and reverse primer containing SalI site, and inserted into these restriction enzyme sites. Following primer sequences were used to amplify 2,396 bp upstream of *BFT* (AT5G62040), 5’-<u>CCTGCAGG</u>GACAGAGTAAATTCAACCACAGCAGGT-3’ and 5’-<u>GTCGAC</u>TTTTCTTTGCTCCAATGTGTTTGCGTTTG-3’; 2,521 bp upstream of AT1G24575, 5’-<u>CCTGCAGG</u>CTCTCAGATCACCGTAAGGGCATAATTATATTTAGGTTCAC-3’ and 5’-<u>GTCGAC</u>GTGATGAGATTTGTGACTGGAGGAGTTTCCAAGTACCATTCTT-3’; 2,488 bp upstream of AT1G67865, 5’-<u>CCTGCAGG</u>ACTTCACATTCTTGGATTCCGTTTGTAATAACTAATGTTTT-3’ and 5’-<u>GTCGAC</u>CCCTCCGGCAACCCCAATAATAAGCTTATCAAGCATTTTTCTT-3’; 1,939 bp upstream of *ROXY10* (AT5G18600), 5’-<u>CCTGCAGG</u>GCAATGGACCGTACGTCTAGGTCACGCATCTTATCCGACAT-3’ and 5’-<u>GTCGAC</u>CACCGGTCTCTCCATCACCATCTTCGTTATCATATCCATTGCT-3’; 2,226 bp upstream of AT2G26695, 5’-<u>CCTGCAGG</u>AATGTAATGTATAATGTGTTCATAAACAGCACCAACTACCC-3’ and 5’-<u>GTCGAC</u>TGCGTGCTGACACGCACCACATAGCCAATCTCCTCCGGTCCAG-3’; 1,911 bp upstream of *PARCL* (AT1G64370), 5’-<u>CCTGCAGG</u>GCCCATCTAATTCCCATTTTAGATGCATGAGTTCAACGCTA-3’ and 5’-<u>GTCGAC</u>GCCTTGAGCCACCTCGTAGTAGTCTTTCTCACGGTTTTCGTAG-3’; 2,719 bp upstream of *ROXY7* (AT2G30540), 5’-<u>CCTGCAGG</u>ATCACCGGTAAGTGACAAGAGAATTGA-3’ and 5’-<u>GTCGAC</u>GGTTTCTTGAAGGAGGTCTCGATCAATCT-3’; 2,574 bp upstream of *CYSTM12* (AT5G04080),5’-<u>CCTGCAGG</u>GAGAATTTGAAGGAGGCTTTGCGTTTTATCTGCTCATCTAA-3’ and 5’-<u>GTCGAC</u>TTGTGGAGGATTTTGATCTCTCATGTCCTGCATCTTCTCAAAA-3’. These constructs were transformed into the *pFT:NTF* plants.

To overexpress cluster 7 highly expressed genes, we amplified coding sequences using the following primer sets. *BFT*, 5’-CACCATGTCAAGAGAAATAGAGCCAC-3’ and 5’-AGTTAATAAGAAGGACGTCGTCG-3’; AT1G24575, 5’-CACCATGGTACTTGGAAACTCCTCCAGTCAC-3’ and 5’-GACTAGGCGCTCTTAGTCATCCAC-3’; AT1G67865, 5’-CACCATGCTTGATAAGCTTATTATTGGGGTTGCC-3’ and 5’-CTTTACTCCCTGTCTTTCTGGCG-3’; *ROXY10*, 5’-CACCATGGATATGATAACGAAGATGGTGATGGAG-3’ and 5’-AGTCAAACCCACAATGCACCAG-3’; AT2G26695, 5’-CACCATGAGCTGGACCGGAGGAG-3’ and 5’-ATTTAGACGCCACCATAATCTCTTG-3’, *SAQR* (AT1G64360), 5’-CACCATGTCGTTTAGAAAAGTAGAGAAGAAACC-3’ and 5’-GATTAGTAATTAGGGAAGTGTTTGCGGC-3’; *RL4* (AT2G18328), 5’-CACCATGGCTTCTAGTTCAATGAGCACC-3’ and 5’-CTTCAATTAGTGTTACGGTACCTAGG-3’; AT1G54575, 5’-CACCATGGTGGATCATCATCTCAAAGC-3’ and 5’-AATTAATTATTCTTTTGTGGCTTGG-3’. Amplified cDNA was cloned into pENTR/D-topo, followed by GATEWAY cloning using pH7SUC2, a binary vector carrying 0.9 kb *SUC2* promoter (15). These vectors were transformed into wild-type plants possessing *pFT:GUS* reporter gene (11).

For gene expression measurements and confocal imaging, surface sterilized seeds were sawn on the 1x Linsmaier and Skoog (LS) media plates without sucrose containing 0.8% (w/v) agar and stratified for more than 2 days at 4 °C before being transferred to the incubator. Plants were grown under LD+FR conditions (100 µmol photons m^-2^ s^-1^, red/far-red ratio = 1.0) for two weeks as described previously (6). For gene expression analysis under –N conditions, plants were grown on normal 1x Murashige and Skoog (MS) medium plates with full nitrogen (+N) for 10 days and transplanted to +N and –N medium plates and grown for 4 additional days. The ion equilibrium of the medium between +N and –N was ensured by replacing KNO_3_ (18.79 mM) and NH_4_NO_3_ (20.61 mM) by KCl (18.79 mM) and NaCl (20.61 mM). For flowering time measurements using T_1_ transgenic lines (Fig. 4B), T_1_ plants were grown on hygromycin selection plates for 10 days and transferred to Sunshine Mix 4 soils (Sun Gro Horticulture). Soils were supplemented with a slow-release fertilizer (Osmocote 14-14-14, Scotts Miracle-Gro) and a pesticide (Bonide, Systemic Granules) and filled in standard flats with inserts (STF-1020-OPEN and STI-0804, T.O. Plastics). For the same experiment using *pSUC2:NIGT1* T_3_ plants (Fig. 5B), surface sterilized seeds were directly sawn on soils and kept in 4 °C for at least 2 days. Plants on soils were kept grown under LD+FR conditions as described previously (6). Tissue clearing and confocal imaging were conducted as previously described (6).

### Nuclei isolation by FANS

Nuclei labeled with NTF proteins were isolated with the modified method described by Galbraith *et al.* (54). Cotyledons and true leaves were detached from 2-week-old plants grown under LD+FR conditions at ZT4 (approximately 100–200 seedlings were used at each sample), and placed on a mixture of 2 mL filter sterilized chopping buffer (20 mM MOPS, 30 mM sodium citrate and 45 mM MgCl_2_) and 20 µL SUPERase-In RNase inhibitor (20 U /µL) (Thermo Fisher Scientific) (**Fig. S1B and C**). We did not use fixing reagents such as formaldehyde in the chopping buffer because it caused clumps of nuclei. Leaves were chopped with the razor blade in 4 °C room until finely crushed in the buffer solution, followed by the addition of 3 mL more chopping buffer and filtration with 30-µm CellTrics filter (Sysmex CORP.). It usually takes approximately 30 min from start chopping leaves to applying the samples for sorting. The filtrated sample solution was directly applied to the sorting with SH800S Cell Sorter equipped with a 100-μm flow tip (SONY). As a sheath solution, we used 0.9% NaCl but not PBS to prevent calcium precipitation by phosphate contained in PBS, which severely decreases the integrity of nuclei. DEPC-treated water was used for all buffers and solutions in this experiment to prevent RNA degradation.

Nuclei were excited by 488 nm collinear laser, and FITC and mCherry channels were used to detect GFP signal and autofluorescence, respectively. The area of GFP-positive nuclei was determined using the parental line *pACT2:BirA* (**Fig. S2**).

### SMART-seq2 library preparation and analysis

For bulked nuclei RNA-seq, approximately 15,000 nuclei were collected from cotyledons and true leaves of *p35S:NTF*, *pCAB2:NTF*, and *SUC2:NTF* lines, while 10,000 (cotyledons) and 3,000 (true leaves) nuclei were collected from *pFT:NTF* due to fewer population of GFP-positive nuclei. Sorted nuclei were directly collected into Buffer RTL (Qiagen), and RNA was extracted according to the manufacture protocol of RNeasy Micro Kit (Qiagen). Three independent biological replicates were produced for cotyledon and true leaf of all transgenic lines.

After RNA integrity was confirmed using High Sensitivity RNA Screen Tape (Agilent), SMART-seq2 libraries were generated according to the manufacturer’s protocol (Clontech). Sequencing was performed on the Illumina NovaSeq 6000 platform through a private sequencing service (Novogene). Since reverse reads were undesirably trimmed due to the presence of in-line indexes in SMART-seq libraries, we used only forward reads of paired-end reads. Approximately 10.8 to 34.6 million forward reads (average, 20.2 million reads) were produced from each sample. By STAR software (version 2.7.6a) (55), reads were mapped to *Arabidopsis* genome sequences from The Arabidopsis Information Resource (TAIR, version 10) (http://www.arabidopsis.org/). DEGs were selected based on the adjusted *P*-value calculated using DEseq2 in the R environment.

### Single-nucleus RNA-seq library preparation

For snRNA-seq experiments, sorted nuclei from true leaves were collected in the mixture of 100 µL 1x PBS and 10 µL SUPERase-In RNase inhibitor in a 15 mL corning tube. Prior to nuclei collection, the corning tube was coated by rotating with 10 mL 1x PBS containing 1% BSA at room temperature to prevent static electricity in the wall. After more than 10,000 nuclei were collected in the corning tube (sorting approximately for 1 hour), 10 µL of 1 mg/mL DAPI solution and 1/100 volume of 20 mg/mL glycogen (Thermo Scientific) were added and centrifugated in 1,000xg for 15 min at 4 °C. Nuclei numbers were determined using a hemocytometer after resuspension with a small volume of 1x PBS. The total number of nuclei varied depending on the plant line and trial but was no more than 11,000.

snRNA-seq was performed using the 10x Single-cell RNA-Seq platform, the Chromium Single Cell Gene Expression Solution (10x Genomics). Two biological replicates of *SUC2:NTF* and one replicate of *pFT:NTF* were produced for a total of three samples.

### Estimating gene expression in individual nucleus

Sequencing of snRNA-seq reads was performed on the Illumina Nextseq 550 platform, followed by the mapping to the TAIR10 Arabidopsis genome using Cellranger version 3.0.1 software.

The Seurat R package (version 4.0.5) (56, 57) was used for the dimensional reduction of our SnRNA-seq data. To remove potential doublets and background, we first filtered out nuclei with less than 100 and more than 5,500 detected genes, and more than 20,000 UMI reads. For UMAP data visualization and cell clustering, all biological replicates were combined, and twenty principal components were compressed using a resolution value of 0.5.

Significantly highly expressed genes in each cluster compared with entire nuclei population were identified using FindMarkers in Seurat with default parameters.

To show the enrichments of specific sets of genes, average expression of ATP biosynthesis, JA responsive, aquaporin, bundle sheath and phloem parenchyma marker genes were visualized using AddModuleScore in Seurat. For bundle sheath and phloem parenchyma markers, we leveraged the top 50 most highly expressed genes in these cell types in previous protoplast-based ScRNA-seq data (16).

### Protein amino acid length

Amino acid length of proteins encoded by genes highly expressed in clusters 4, 5, and 7 were obtained from the Bulk Data Retrieval tool in TAIR database (https://www.arabidopsis.org/tools/bulk/sequences/index.jsp).

### Promoter *cis*-enrichment analysis

To identify highly enriched *cis-*elements in cluster 7 highly expressed genes, promoter sequences of 268 genes highly expressed in cluster 7 and randomly selected 3,000 genes by R coding were extracted using the Eukaryotic Promoter Database (37, 38). Obtained promoter sequences were next submitted to Simple Enrichment Analysis (SEA) (58) in MEME Suite server (version 5.4.1) to elucidate what *cis*-elements are enriched in cluster 7 highly expressed genes through a comparison with random 3,000 genes.

### Yeast one-hybrid screening

Four tandem repeats of S1/S2 elements of *FT* promoter that include a CO-responsive (CORE) element (41, 42) were generated through restriction enzyme-mediated ligation. Forward and reverse oligonucleotides containing S1/S2 element with HindIII (CORE-F1: 5’-<u>AGCTT</u>ACTGTGTGATGTACGTAGAATCAGTTTTAGATTCTAGTA CTGTGTGATGTACGTAGAATCAGTTTTAGATTCTA<u>G</u>-3’), EcoRI (CORE-R1: 5’-<u>AATTC</u>CTAGAATCTAAAACTGATTCTACGTACATCACACAGTACTAGAATCTAAAACTGATTCT ACGTACATCACACAGT<u>A</u>-3’), SacI (CORE-F2: 5’-<u>C</u>ACTGTGTGATGTACGTAGAATCAGTTTTAGATTCTAGTACTGTGTGATGTAC GTAGAATCAGTTTTAGATTCTAG<u>GGTAC</u>-3’), and KpnI (CORE-R2: 5’-<u>C</u>CTAGAATCTAAAACTGATTCTACGTACATCACACAGTACTAGAATCTAAAACTGATTCTACG

TACATCACACAGT<u>GAGCT</u>-3’) recognition sequences were chemically synthesized from Genewiz (https://www.genewiz.com/). The oligonucleotides complementary to each other were denatured for 10 min at 95 LC, then annealed for 30 min at RT. The resultant product was inserted into pENTR/D-TOPO vector harboring MCS sites (HindIII, EcoRI, SacI and KpnI) through restriction enzyme-mediated ligation. Four tandem S1/S2 elements in pENTR/D-TOPO were then inserted into pY1-gLUC59-GW vector (43) using LR clonase II (Invitrogen). The resultant pY1-gLUC59-GW-2XCORE plasmids were used for Y1H screening analysis.

### qRT-PCR analysis using total RNA

Total RNA extraction, cDNA synthesis and qRT-PCR were performed as previously described (6).

*ISOPENTENYL PYROPHOSPHATE / DIMETHYLALLYL PYROPHOSPHATE ISOMERASE* (*IPP2*) and *PROTEIN PHOSPHATASE 2A SUBUNIT A3* (*PP2AA3*) were used as reference genes. For statistical tests, relative expression levels were log_2_-transformed to meet the requirements for homogeneity of variance. qPCR primers used in this study are listed in Table S2.

### RNA-seq analysis using total RNA

Two-week-old seedlings of *pSUC2:NIGT1.2* and *NIGT1.4* grown on 1xLS media plates under LD+FR conditions were harvested at ZT4. RNA extraction and RNA-seq library preparation were conducted as described previously (15, 59). As a reference, our previous gene expression data of wild-type plants grown at the exact same conditions was compared with *pSUC2:NIGT1* lines(15).

### Tobacco transient promoter LUC assay

We amplified 1 kb upstream of *FT* (*pFT*) using the primer sets, pFT-FW (5’-CACCATAATATGGCCGCTTGTTTATA-3’) and pFT-RE (5’-CTTTGATCTTGAACAAACAGGTGG-3’), and cloned it into pENTR/D-topo. To mutate 2 of 3 potential NIGT1-binding sites, we synthesized the Megaprimer #1 by PCR using the forward primer pFTmut1 (5’-CAATGTGTGATGTACGTATACTCAGTTTTAGAGTATAGTACATCAATAGACAAGAAAAAG-3’) and pFT-RE. Next, to mutate the rest of NIGT1-bind site, the Megaprimer #2 was synthesized using the forward primer pFTmut2 (5’-CTACCAAGTGGGAGATATAATTTGAATTAATTCCAGTGTATTAGTGTGGTG) and Megaprimer #1. Subsequently, we amplified 1 kb promoter *FT* with mutations in all 3 NIGT1-bindning sites (*pFTm*), we amplified DNA using pFT-FW and Megaprimer #2, and cloned it into pENTR/D-topo. Finally, *pFT* and *pFTm* regions were cloned into pFLASH, the binary GATEWAY vector for promotor LUC assay (60).

The promoter LUC assay in *N. benthamiana* leaf was conducted as previously described with slight modifications (61). *Agrobacterium* carrying binary vectors was cultured in LB media containing 20 µM acetosyringone and antibiotics overnight and subsequently precipitated at 4,000 rpm at room temperature for 15 min. The *Agrobacterium* pellet was resuspended using inoculation buffer (10 mM MgCl_2_, 10 mM MES-KOH pH5.6, and 100 µM acetosyringone). After incubation for 3 hours at room temperature, OD_600_ was adjusted to 0.2 and infiltrated into the abaxial side of tobacco leaves using a 1 mL syringe. *Agrobacterium* carrying reporters (*pFT:LUC* and *pFTm:LUC*), effector (*p35S:NIGT1.2*, *p35S:NIGT1.4* and *p35S:GFP* in the binary vector pB7WG2) were co-inoculated with pBIN61 P19 (62). Two days after inoculation, leaf discs were punched out and their LUC activities were measured using Dual-Luciferase Reporter Assay System kit (Promega) and SpectraMax M5e (Molecular Devices).

## Data availability

Bulk RNA-seq from sorted nuclei can be found at the NCBI short read archive bio project PRJNA1098062. snRNA-seq can be found at NCBI GEO under GSE273032. The RNA-seq data of *pSUC2:NIGT1.2* and *pSUC2:NIGT1.4* plants have been deposited in the DNA Data Bank of Japan (DDBJ) Sequence Read Archive under PRJDB17784.

## Acknowledgments

We thank N.M. Belliveau and D.W Galbraith for technical advice for FACS, Y. Mizuta for providing research materials, M. Ashikari, H. Tsuji, K. Nagai, and M. Mizutani for scientific discussion. We also thank T. Niwa, A. Furuta, S. Ishikawa and T. Takagi for technical assistance. This research was supported by grants from the National Institutes of Health grant (R01GM079712 to J.T.C., C.Q., and T.I.; NIGMS MIRA grant no. 1R35GM139532 to C.Q.), from the National Science Foundation (PlantSynBio grant no. 2240888 to C.Q), Japan MEXT KAKENHI (JP20H05910 and JP22H04978 to T.I.), JSPS KAKENHI (24K18139 to H.T.) and National Science Foundation grant (NSF-IOS #1755452 to J.L.P.-P.).

## Author Contributions

H.T. and T.I conceptualized the study.

Methodology: H.T. and C.M.A.

Investigation: H.T., S.I, J.S.S, A.K., A.K.H., N.L., T.S., J.W., C.Y., C.T.N., J.L.P-P and T.I.

Formal analysis: H.T., T.S., K.L.B., J.L.P.-P. and J.T.C. Data curation: H.T. T.S. and J.T.C.

Validation: H.T., A.K.H., and T.I. Visualization: H.T.

Funding acquisition: H.T., J.L.P.-P., J.T.C., C.Q. and T.I.

Resources: S.I., D.K., T.K. and J.L.P.-P.

Project administration: H.T., C.Q. and T.I.

Supervision: Y.S., Y.T., J.L.P.-P., C.Q. and T.I.

H.T., S.I, J.S., A.K.H, J.T.C., and C.Q. and T.I.

wrote the manuscript. N.L. reviewed and commented on the manuscript.

## References

1. F. Andres, G. Coupland, The genetic basis of flowering responses to seasonal cues. Nat Rev Genet 13, 627–639 (2012).

2. Y. H. Song, J. S. Shim, H. A. Kinmonth-Schultz, T. Imaizumi, Photoperiodic flowering: time measurement mechanisms in leaves. Annu Rev Plant Biol 66, 441–464 (2015).

3. Y. Zhu, S. Klasfeld, D. Wagner, Molecular regulation of plant developmental transitions and plant architecture via PEPB family proteins: an update on mechanism of action. J Exp Bot 72, 2301–2311 (2021).

4. S. N. Freytes, M. Canelo, P. D. Cerdán, Regulation of Flowering Time: When and Where? Curr Opin Plant Biol 63, 102049 (2021).

5. S. P. Pandey, R. M. Benstein, Y. Wang, M. Schmid, Epigenetic Regulation of Temperature Responses - Past Successes and Future Challenges. J Exp Bot 72, 7482–7497 (2021).

6. H. Takagi, A. K. Hempton, T. Imaizumi, Photoperiodic flowering in Arabidopsis: Multilayered regulatory mechanisms of CONSTANS and the florigen FLOWERING LOCUS T. Plant Commun. 4, 100552 (2023).

7. L. Corbesier et al., FT protein movement contributes to long-distance signaling in floral induction of Arabidopsis. Science 316, 1030–1033 (2007).

8. S. Tamaki, S. Matsuo, H. L. Wong, S. Yokoi, K. Shimamoto, Hd3a protein is a mobile flowering signal in rice. Science 316, 1033–1036 (2007).

9. J. Mathieu, N. Warthmann, F. Küttner, M. Schmid, Export of FT protein from phloem companion cells is sufficient for floral induction in Arabidopsis. Curr Biol 17, 1055–1060 (2007).

10. M. Notaguchi et al., Long-distance, graft-transmissible action of Arabidopsis FLOWERING LOCUS T protein to promote flowering. Plant Cell Physiol 49, 1645–1658 (2008).

11. S. Takada, K. Goto, TERMINAL FLOWER2, an Arabidopsis homolog of HETEROCHROMATIN PROTEIN1, counteracts the activation of FLOWERING LOCUS T by CONSTANS in the vascular tissues of leaves to regulate flowering time. Plant Cell 15, 2856–2865 (2003).

12. Q. Chen et al., FLOWERING LOCUS T mRNA is synthesized in specialized companion cells in Arabidopsis and Maryland Mammoth tobacco leaf veins. Proc. Natl. Acad. Sci. U.S.A. 115, 2830–2835 (2018).

13. Y. H. Song et al., Molecular basis of flowering under natural long-day conditions in Arabidopsis. Nat. Plants 4, 824–835 (2018).

14. N. Lee et al., The FLOWERING LOCUS T gene expression is controlled by high-irradiance response and external coincidence mechanism in long days in Arabidopsis. New Phytol. 239, 208–221 (2023).

15. H. Takagi et al., Florigen-producing cells express FPF1-LIKE PROTEIN 1 to accelerate flowering and stem growth in Arabidopsis. Dev Cell 60, 1822–1837 e1828 (2025).

16. J. Y. Kim et al., Distinct identities of leaf phloem cells revealed by single cell transcriptomics. Plant Cell 33, 511–530 (2021).

17. R. B. Deal, S. Henikoff, The INTACT method for cell type-specific gene expression and chromatin profiling in Arabidopsis thaliana. Nat. Protoc. 6, 56–68 (2011).

18. D. Shi et al., Tissue-specific transcriptome profiling of the Arabidopsis inflorescence stem reveals local cellular signatures. Plant Cell 33, 200–223 (2021).

19. S. Ito et al., FLOWERING BHLH transcriptional activators control expression of the photoperiodic flowering regulator CONSTANS in Arabidopsis. Proc Natl Acad Sci U S A 109, 3582–3587 (2012).

20. Y. Zhou et al., Metascape provides a biologist-oriented resource for the analysis of systems-level datasets. Nat. Commun. 10, 1523 (2019).

21. L. McInnes, J. Healy, J. Melville, UMAP: Uniform manifold approximation and projection for dimension reduction. arXiv preprint arXiv 1802, 03426 (2018).

22. H. Hunt et al., Analysis of companion cell and phloem metabolism using a transcriptome-guided model of Arabidopsis metabolism. Plant Physiol 192, 1359–1377 (2023).

23. S. Kushwah et al., Arabidopsis XTH4 and XTH9 contribute to wood cell expansion and secondary wall formation. Plant Physiol 182, 1946–1965 (2020).

24. C. G. Litholdo, Jr.,et al, Proteomic identification of putative MicroRNA394 target genes in Arabidopsis thaliana identifies major latex protein family members critical for normal development. Mol Cell Proteomics 15, 2033–2047 (2016).

25. H. Wang et al., Regulatory function of Arabidopsis lipid transfer protein 1 (LTP1) in ethylene response and signaling. Plant Mol Biol 91, 471–484 (2016).

26. K. Prado et al., Regulation of Arabidopsis leaf hydraulics involves light-dependent phosphorylation of aquaporins in veins. Plant Cell 25, 1029–1039 (2013).

27. A. Chauvin, A. Lenglet, J. L. Wolfender, E. E. Farmer, Paired hierarchical organization of 13-lipoxygenases in Arabidopsis. Plants 5, 6 (2016).

28. L. Kienow et al., Jasmonates meet fatty acids: functional analysis of a new acyl-coenzyme A synthetase family from Arabidopsis thaliana. J Exp Bot 59, 403–419 (2008).

29. S. Li, J. Ma, P. Liu, OPR3 is expressed in phloem cells and is vital for lateral root development in Arabidopsis. Canadian Journal of Plant Science 93, 165–170 (2013).

30. S. Aubry, R. D. Smith-Unna, C. M. Boursnell, S. Kopriva, J. M. Hibberd, Transcript residency on ribosomes reveals a key role for the Arabidopsis thaliana bundle sheath in sulfur and glucosinolate metabolism. Plant J 78, 659–673 (2014).

31. K. Torii et al., A guiding role of the Arabidopsis circadian clock in cell differentiation revealed by time-series single-cell RNA sequencing. Cell Reports 40, 111059 (2022).

32. M. Biedron, A. Banasiak, Auxin-mediated regulation of vascular patterning in Arabidopsis thaliana leaves. Plant Cell Rep 37, 1215–1229 (2018).

33. Y. Yang, A. Costa, N. Leonhardt, R. S. Siegel, J. I. Schroeder, Isolation of a strong Arabidopsis guard cell promoter and its potential as a research tool. Plant Methods 4, 6 (2008).

34. D. C. Jones et al., A clade-specific Arabidopsis gene connects primary metabolism and senescence. Front Plant Sci 7, 983 (2016).

35. C. E. Baxter, M. M. Costa, E. S. Coen, Diversification and co-option of RAD-like genes in the evolution of floral asymmetry. Plant J 52, 105–113 (2007).

36. J. Y. Ryu et al., The Arabidopsis floral repressor BFT delays flowering by competing with FT for FD binding under high salinity. Mol Plant 7, 377–387 (2014).

37. R. Dreos, G. Ambrosini, R. C. Perier, P. Bucher, The Eukaryotic Promoter Database: expansion of EPDnew and new promoter analysis tools. Nucleic Acids Res 43, D92–96 (2015).

38. P. Meylan, R. Dreos, G. Ambrosini, R. Groux, P. Bucher, EPD in 2020: enhanced data visualization and extension to ncRNA promoters. Nucleic Acids Res 48, D65–D69 (2020).

39. T. Kiba et al., Repression of nitrogen starvation responses by members of the Arabidopsis GARP-type transcription factor NIGT1/HRS1 subfamily. Plant Cell 30, 925–945 (2018).

40. Y. Maeda et al., A NIGT1-centred transcriptional cascade regulates nitrate signalling and incorporates phosphorus starvation signals in Arabidopsis. Nat Commun 9, 1376 (2018).

41. J. Adrian et al., cis-Regulatory elements and chromatin state coordinately control temporal and spatial expression of FLOWERING LOCUS T in Arabidopsis. Plant Cell 22, 1425–1440 (2010).

42. S. B. Tiwari et al., The flowering time regulator CONSTANS is recruited to the FLOWERING LOCUS T promoter via a unique cis-element. New Phytol 187, 57–66 (2010).

43. K. Bonaldi, Z. Li, S. E. Kang, G. Breton, J. L. Pruneda-Paz, Novel cell surface luciferase reporter for high-throughput yeast one-hybrid screens. Nucleic Acids Res 45, e157 (2017).

44. Z. Li, K. Bonaldi, S. E. Kang, J. L. Pruneda-Paz, High-Throughput Yeast One-Hybrid Screens Using a Cell Surface gLUC Reporter. Curr Protoc Plant Biol 4, e20086 (2019).

45. M. Sanagi et al., Low nitrogen conditions accelerate flowering by modulating the phosphorylation state of FLOWERING BHLH 4 in Arabidopsis. Proc Natl Acad Sci U S A 118, e2022942118 (2021).

46. S. J. Yoo, et al., BROTHER OF FT AND TFL1 (BFT) has TFL1-like activity and functions redundantly with TFL1 in inflorescence meristem development in Arabidopsis. Plant J 63, 241–253 (2010).

47. Y. Higuchi, Florigen and anti-florigen: flowering regulation in horticultural crops. Breed Sci 68, 109–118 (2018).

48. Y. Higuchi et al., The gated induction system of a systemic floral inhibitor, antiflorigen, determines obligate short-day flowering in chrysanthemums. Proc Natl Acad Sci U S A 110, 17137–17142 (2013).

49. S. Soyk et al., Variation in the flowering gene SELF PRUNING 5G promotes day-neutrality and early yield in tomato. Nat Genet 49, 162–168 (2017).

50. Y. Zhu et al., TERMINAL FLOWER 1-FD complex target genes and competition with FLOWERING LOCUS T. Nat Commun 11, 5118 (2020).

51. S. M. McKim, Moving on up - controlling internode growth. New Phytol. 226, 672–678 (2020).

52. B. B. Maxwell, C. R. Andersson, D. S. Poole, S. A. Kay, J. Chory, HY5, Circadian Clock-Associated 1, and a cis-element, DET1 dark response element, mediate DET1 regulation of chlorophyll a/b-binding protein 2 expression. Plant Physiol 133, 1565–1577 (2003).

53. T. Nakagawa et al., Development of R4 gateway binary vectors (R4pGWB) enabling high-throughput promoter swapping for plant research. Biosci Biotechnol Biochem 72, 624–629 (2008).

54. D. W. Galbraith, “Flow Cytometry and Sorting in Arabidopsis” in Arabidopsis Protocols, 3rd Edition, J. J. SanchezSerrano, J. Salinas, Eds. (2014), vol. 1062, pp. 509–537.

55. A. Dobin et al., STAR: ultrafast universal RNA-seq aligner. Bioinformatics 29, 15–21 (2013).

56. Y. Hao et al., Integrated analysis of multimodal single-cell data. Cell 184, 3573–3587 e3529 (2021).

57. R. Satija, J. A. Farrell, D. Gennert, A. F. Schier, A. Regev, Spatial reconstruction of single-cell gene expression data. Nat Biotechnol 33, 495–502 (2015).

58. T. L. Bailey, C. E. Grant, SEA: Simple Enrichment Analysis of motifs. bioRxiv 10.1101/2021.08.23.457422, 2021.2008.2023.457422 (2021).

59. T. Suzuki et al., The DROL1 subunit of U5 snRNP in the spliceosome is specifically required to splice AT-AC-type introns in Arabidopsis. Plant J. 109, 633–648 (2022).

60. T. F. Schultz, T. Kiyosue, M. Yanovsky, M. Wada, S. A. Kay, A role for LKP2 in the circadian clock of Arabidopsis. Plant Cell 13, 2659–2670 (2001).

61. A. Kubota et al., TCP4-dependent induction of CONSTANS transcription requires GIGANTEA in photoperiodic flowering in Arabidopsis. PLoS Genet. 13, e1006856 (2017).

62. O. Voinnet, S. Rivas, P. Mestre, D. Baulcombe, An enhanced transient expression system in plants based on suppression of gene silencing by the p19 protein of tomato bushy stunt virus. Plant J 33, 949–956 (2003).

