## Supplemental information for "Companion cells with high florigen production express other small proteins and reveal a nitrogen-sensitive *FT* repressor"

**This PDF file includes:**

Figures S1 to S17

Table S1 and S2

Figure S1 to S17

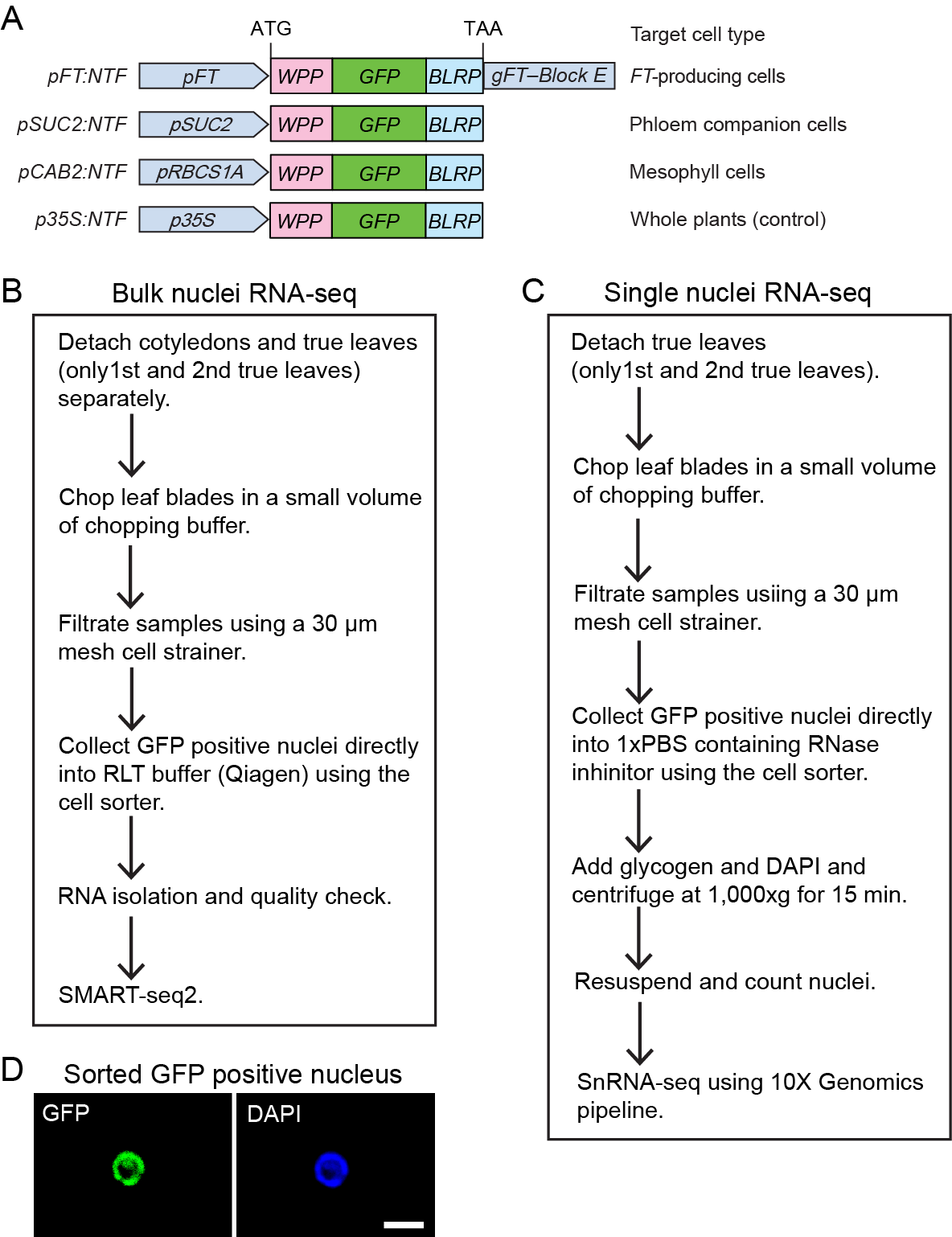

**Supplemental Figure S1.** (A) A schematic diagram of the constructs containing tissue-specific promoters and nuclear targeting fusion protein (NTF). NTF possesses WPP domain, nuclear envelope-targeting domain; GFP, green fluorescent protein; and BLRP, biotin ligase recognition peptides. (B and C) Procedures for preparing sorted nuclei for bulk RNA-seq (A) and single nuclei RNA-seq (B). (C) An example of a sorted GFP-positive nucleus. DAPI stain of the same nucleus is shown. Scale bar indicates 10 µm.

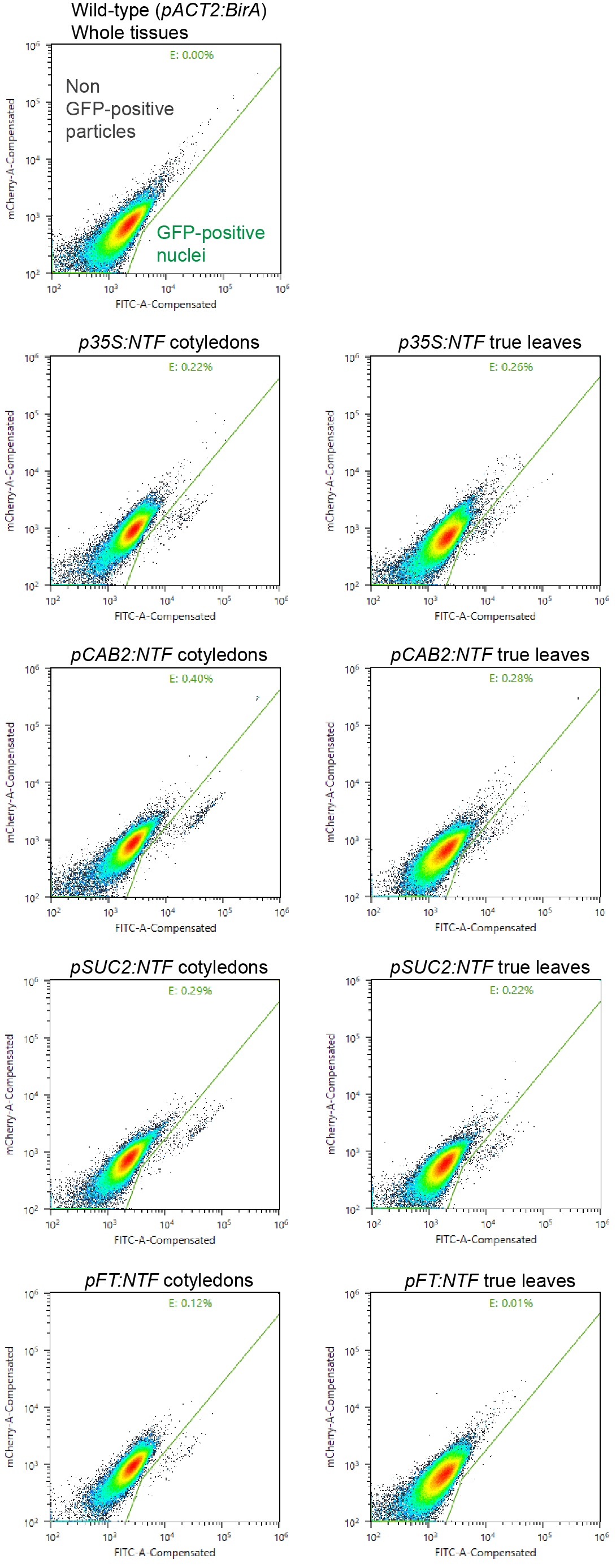

**Supplemental Figure S2.** Plots showing 100,000 events of fluorescence-activated nuclei sorting (FANS). FITC and mCherry channels detect GFP and autofluorescence, respectively. Areas for GFP-positive nuclei and negative particles were determined using whole tissues of the genetic background line *pACT2:BirA*. GFP-positive nuclei were obtained from cotyledons and true leaves of all transgenic lines.

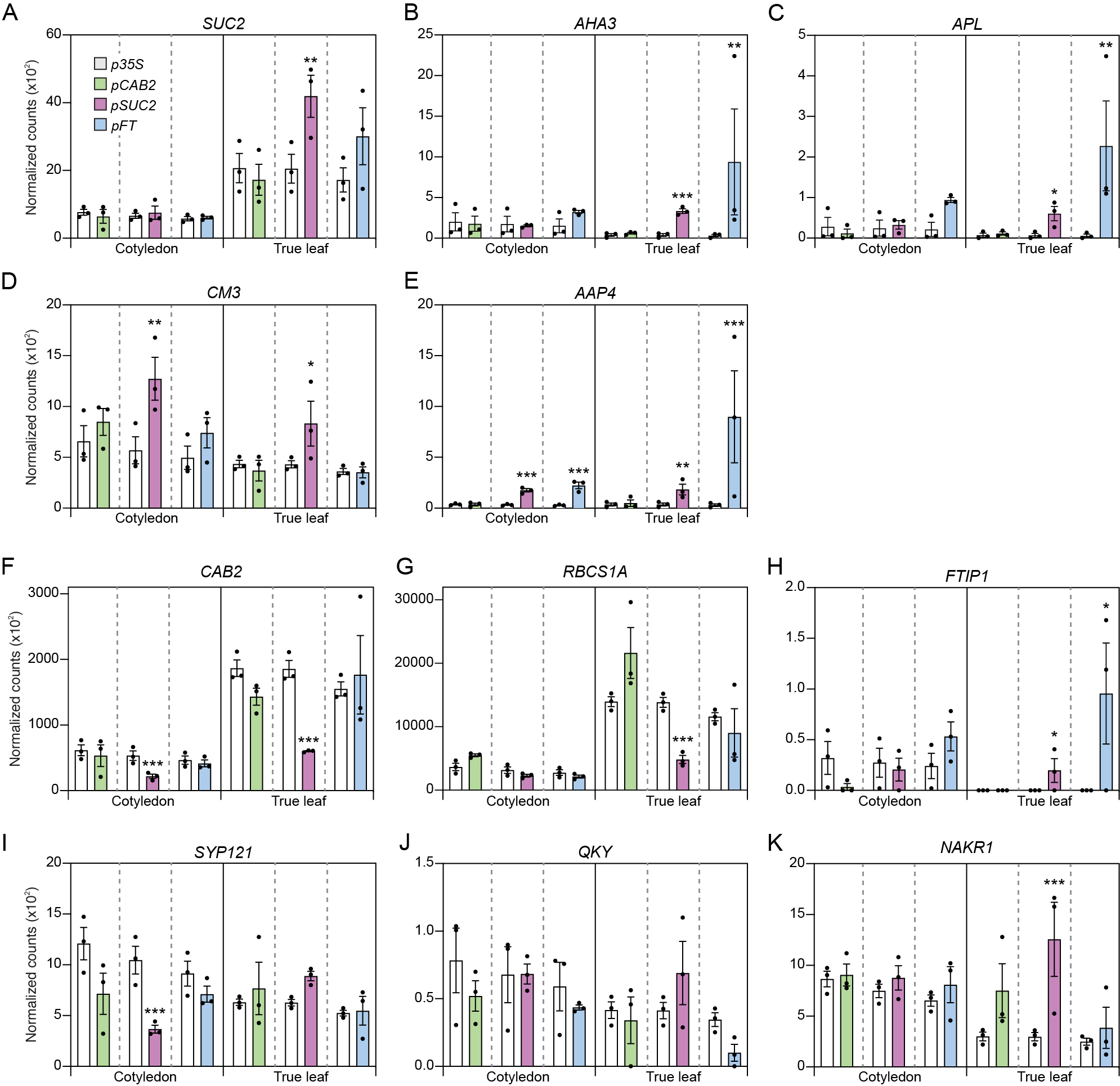

**Supplemental Figure S3.** DEseq2-normalized expression of phloem companion cell marker genes (A–E), mesophyll cell marker genes (F and G), and FT transporting genes (H–I) from sorted nuclei bulk RNA-seq of cotyledons and true leaves. Asterisks denote significant differences from *p35S:NTF* line (**padj*<0.05, ***padj*<0.01, ****padj*<0.001).

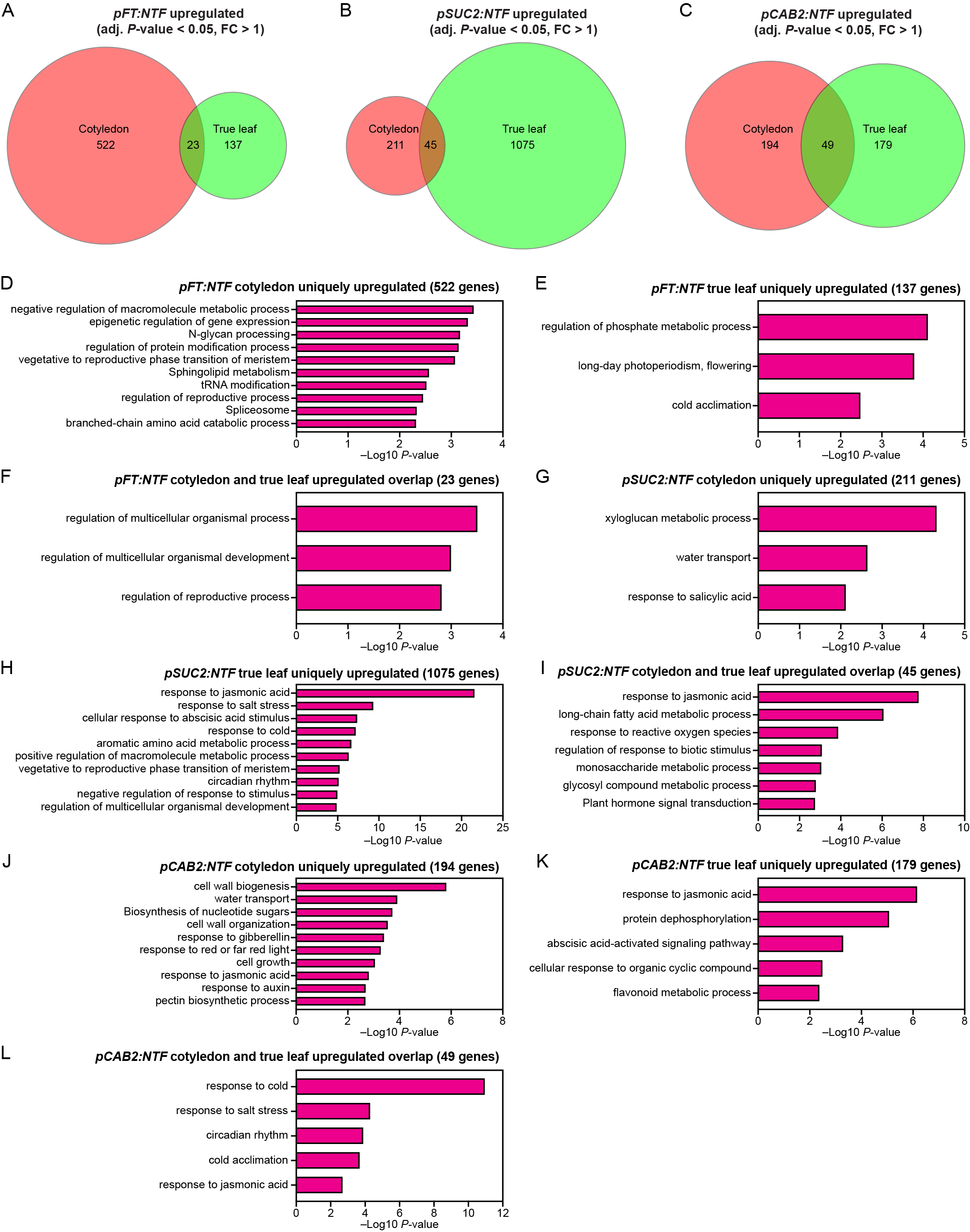

**Supplemental Figure S4.** Cotyledons and true leaves express unique sets of upregulated genes in the sorted nuclei. (A–C) Quantitative Venn diagrams display the overlap between significantly upregulated genes from cotyledons and true leaves of *pFT:NTF* (A), *pSUC2:NTF* (B), and *pCAB2:NTF* lines (C) compared with the *p35S:NTF* line. (D–L) The top 10 Metascape enriched terms from genes uniquely upregulated in cotyledons (D, G, and J), true leaves (E, H, and K), and both (F, I, and L) of *pFT:NTF* (D–F), *pSUC2:NTF* (G–I), and *pCAB2:NTF* (J–L).

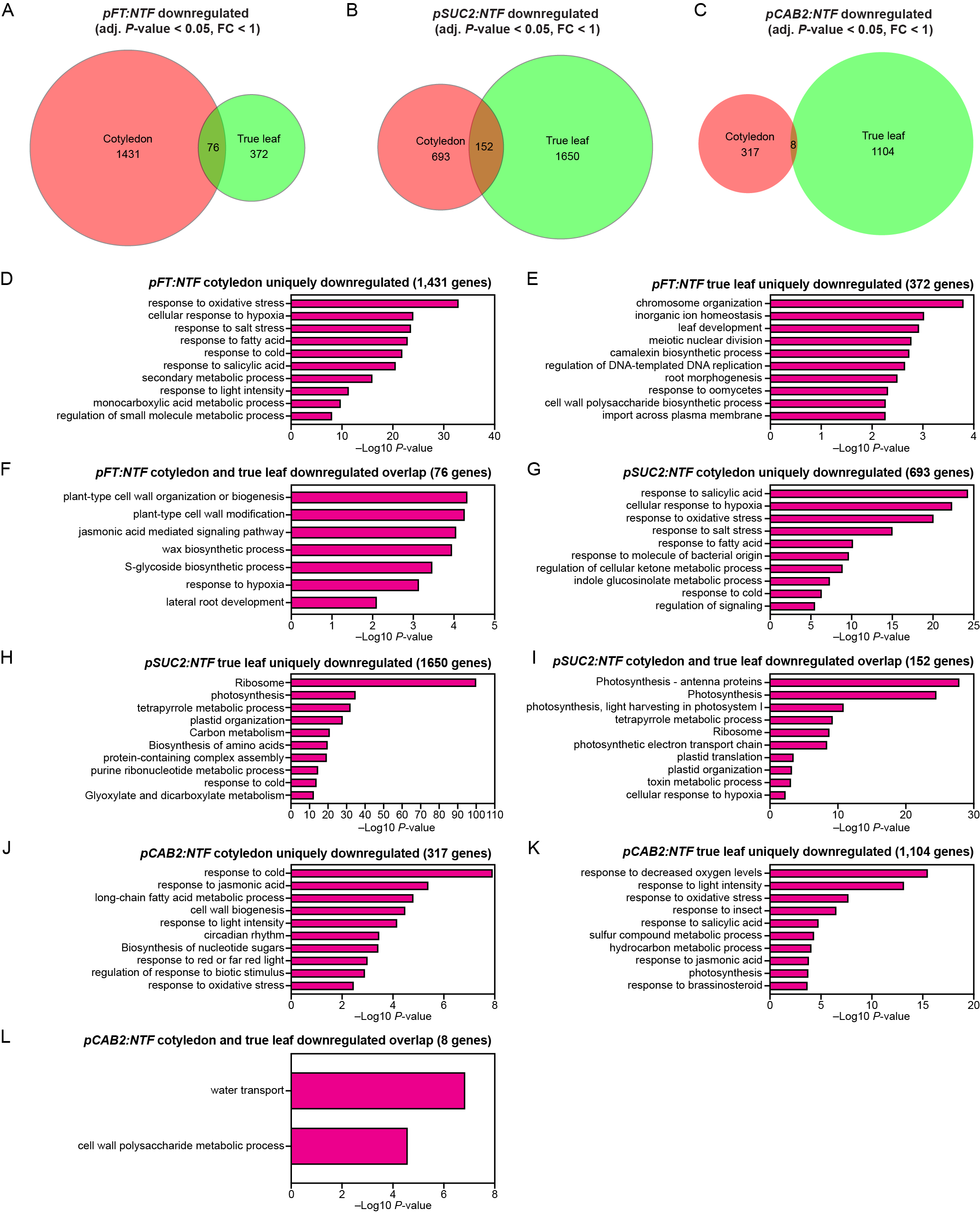

**Supplemental Figure S5.** Cotyledons and true leaves express unique sets of downregulated genes in the sorted nuclei. (A–C) Quantitative Venn diagrams display the overlap between significantly downregulated genes from cotyledons and true leaves of *pFT:NTF* (A), *pSUC2:NTF* (B) and *pCAB2:NTF* lines (C) compared with the *p35S:NTF* line. (D–L) The top 10 Metascape enriched terms from genes uniquely downregulated in cotyledons (D, G, and J), true leaves (E, H, and K), and both (F, I, and L) of *pFT:NTF* (D–F), *pSUC2:NTF* (G–I), and *pCAB2:NTF* (J–L).

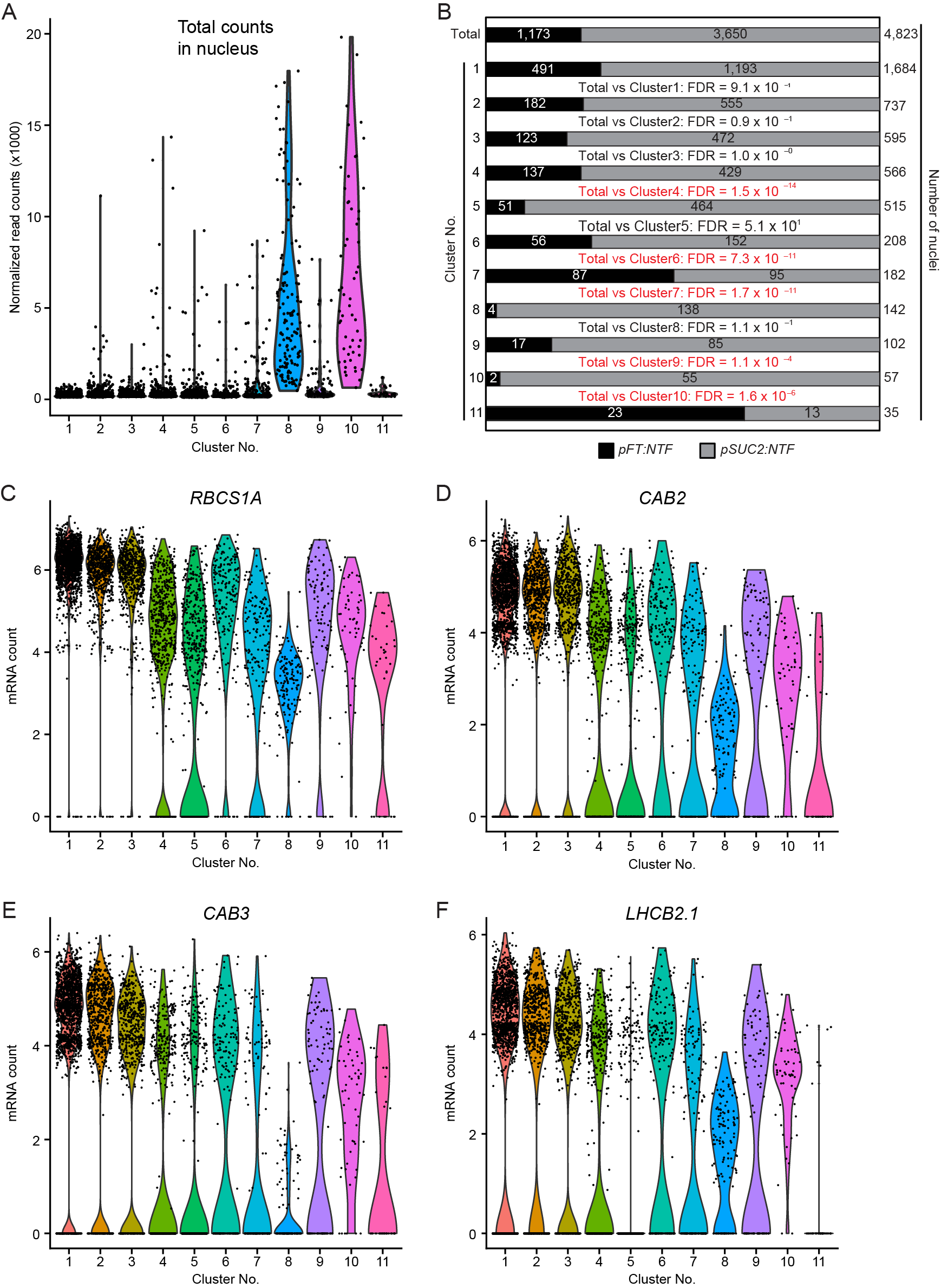

**Supplemental Figure S6.** Characteristics of UMAP clusters 8 and 10. (A) Total read counts in individual nuclei in clusters 8 and 10. (B) Proportions of nuclei from each sample in clusters 8 and 10. The proportions of *pFT:NTF-* and *pSUC2:NTF*-derived nuclei in total and in each cluster were compared using Fisher’s exact test. Clusters significantly different from the total population were denoted with red letters. (C–F) Violin plots showing expression of mesophyll cell marker genes in clusters 8 and 10.

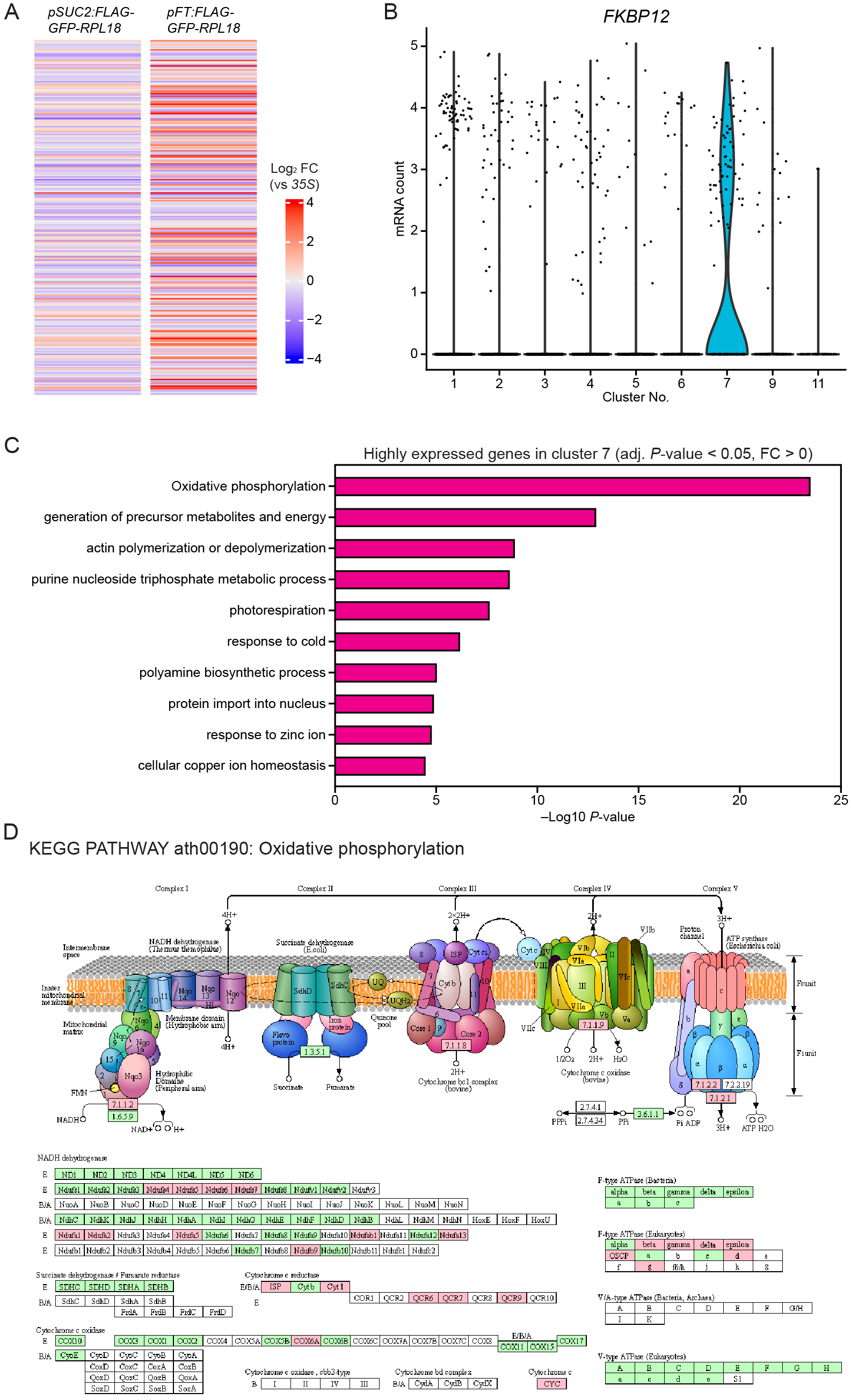

**Supplemental Figure S7.** Characteristics of the *FT*-expressing cluster 7*.* (A) Expression of the 268 genes differentially expressed in cluster 7 in the previous translatome analysis in *SUC2* and *FT* expressing cells using whole plants of *pSUC2:FLAG-GFP-RPL18* and *pFT:FLAG-GFP-RPL18* lines. The color indicates log2 fold-change (FC) compared to the *p35S:FLAG-GFP-RPL18* data. (B) Violin plots showing expression of CO stabilizing gene *FKBP12* in cluster 7. (C) Top 10 Metascape terms enriched in clusters 7. (D) Oxidative phosphorylation pathway (ath00190) showing genes involved in proton and ATP synthesis. Genes highlighted in pink are significantly enriched in cluster 7, while those denoted in green are not enriched in cluster 7. Genes without color are not *Arabidopsis* genes.

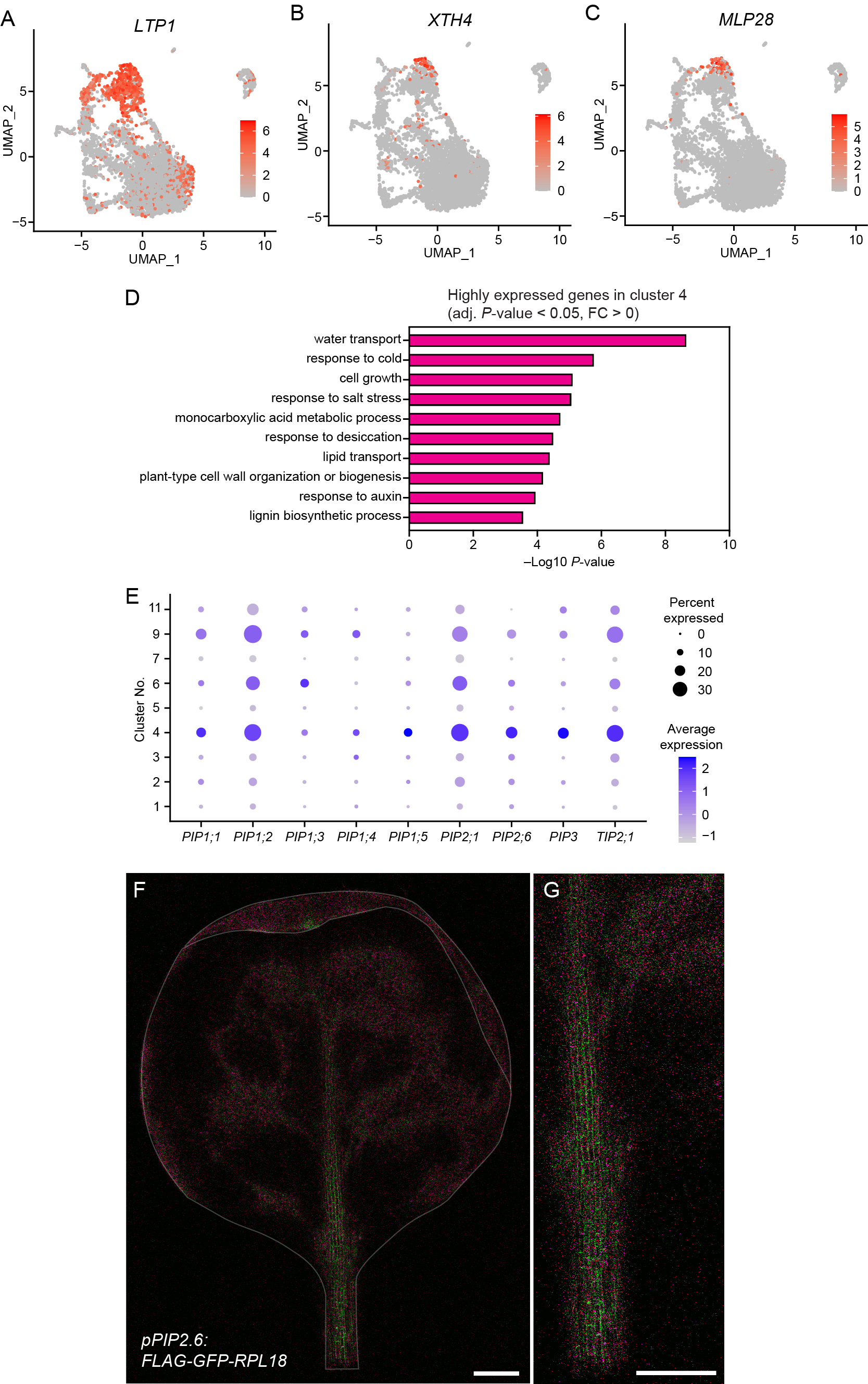

**Supplemental Figure S8.** Characteristics of UMAP cluster 4. (A–C) Leaf vasculature-specific genes. (D) Top 10 Metascape terms enriched in clusters 4. (E) Dot plot showing expression levels of genes encoding aquaporins in each cell cluster. Sizes and colors of dots indicate percents of nuclei expressing genes and average expression levels, respectively. (F) Spatial promoter activity of *PIP2.6* marked with *FLAG-GFP-RPL18* in 2-week-old true leaf. Abaxial side of leaf was imaged. The image is generated by the spectrum imaging capturing GFP (green) and autofluorescence spectrum (magenta). The outline of the leaf is indicated by a white thin line (note the tip of the leaf was curled). (G) An enlarged image of a part of the image (F) that shows the strong promoter activity of *PIP2.6* in the main vein. Scale bar, 500 µm.

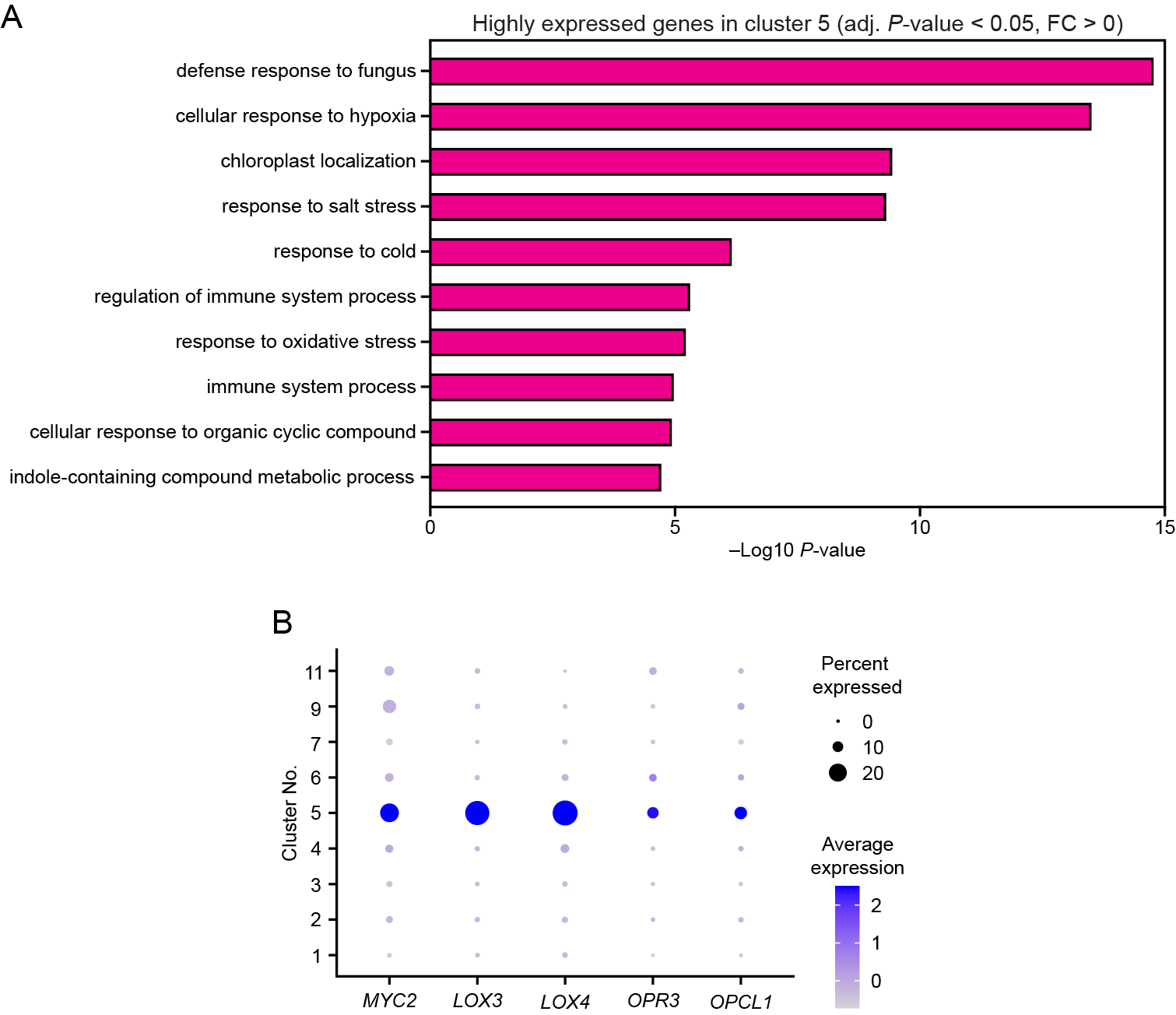

**Supplemental Figure S9.** Characteristics of cluster 5. (A) Top 10 Metascape terms enriched in cluster 5. (B) Dot plot showing expression levels of genes encoding JA-biosynthetic and JA-responsive genes in each cell cluster. Sizes and colors of dots indicate percents of nuclei expressing these genes and average expression levels, respectively.

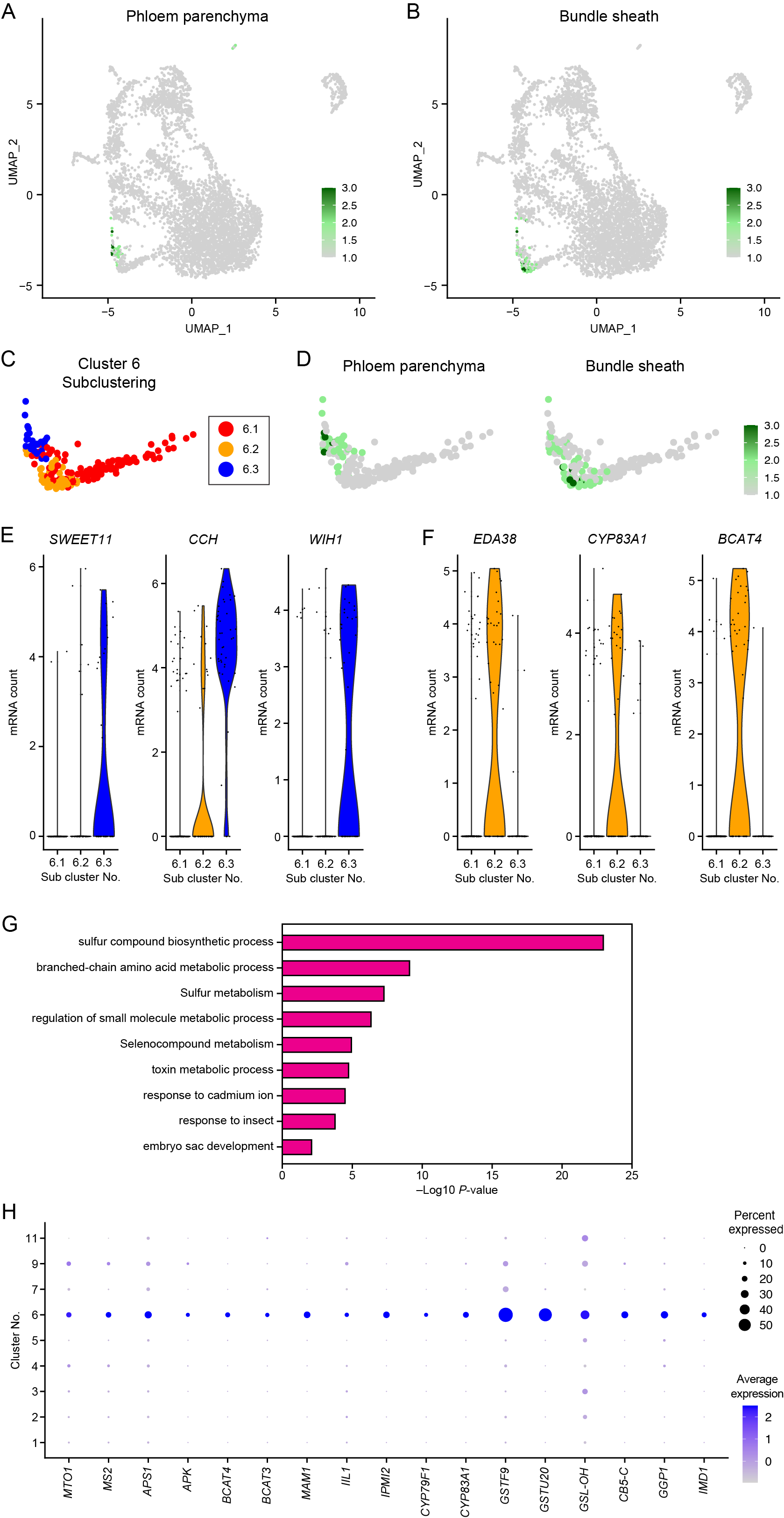

**Supplemental Figure S10.** Characteristics of cluster 6. (A and B) Average expression of top 50 phloem parenchyma (A) and bundle sheath marker genes (B) (1). (C) Subclustering of cluster 6. Colors indicate the positions of each subcluster. (D) Phloem parenchyma marker genes are enriched in subcluster 6.3 and bundle sheath marker genes in subcluster 6.2. (E and F) Violin plots showing expression of phloem parenchyma (E) and bundle sheath marker genes (F). (G) Top 10 Metascape terms enriched in clusters 6. (H) Dot plots showing expression levels of genes related to glucosinolate biosynthesis in each cell cluster. Sizes and colors of dots indicate percents of nuclei expressing genes and average expression levels, respectively.

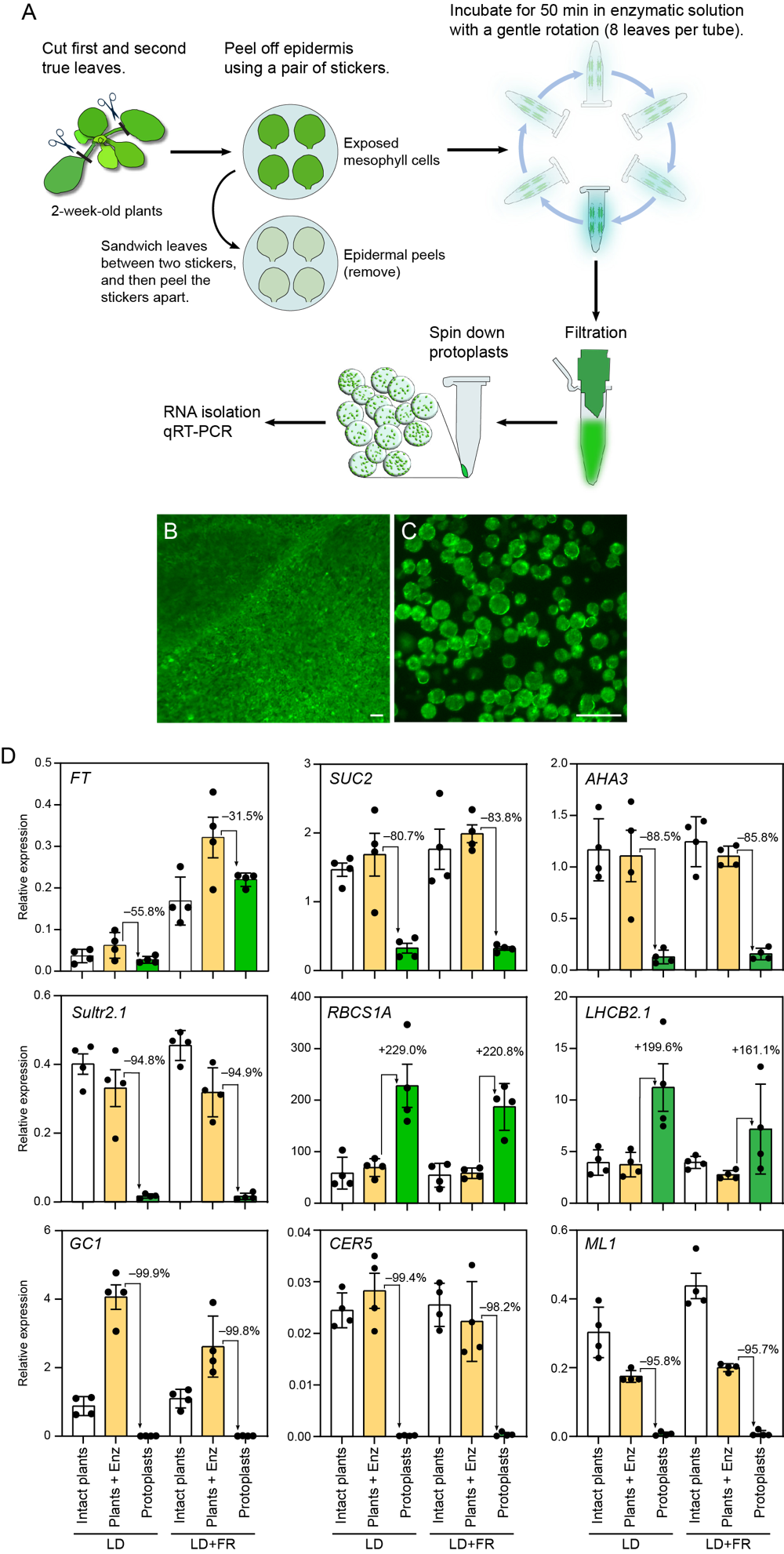

**Supplemental Figure S11.** Protoplast isolation procedure and the expression of tissue-specific marker genes in isolated protoplasts. (A) A schematic diagram of protoplast isolation from true leaves. (B–C) Mesophyll-specific GFP expression of the intact leaf (B) and GFP fluorescence of isolated protoplasts (C) derived from leaves of the *pRBCS1A:FLAG-GFP-RPL18* line. Scale bars, 50 µm. (D) Gene expression analysis using quantitative RT-PCR in mesophyll-cell protoplasts. Intact plants and Plants + Enz indicate detached true leaves with and without protoplasting enzymatic treatment. Epidermis was removed using adhesive tape prior to enzyme treatment but not for Plants + Enz. The results are means ± SEM with each dot representing biological replicates (*n* = 4). Percentage values indicate relative differences.

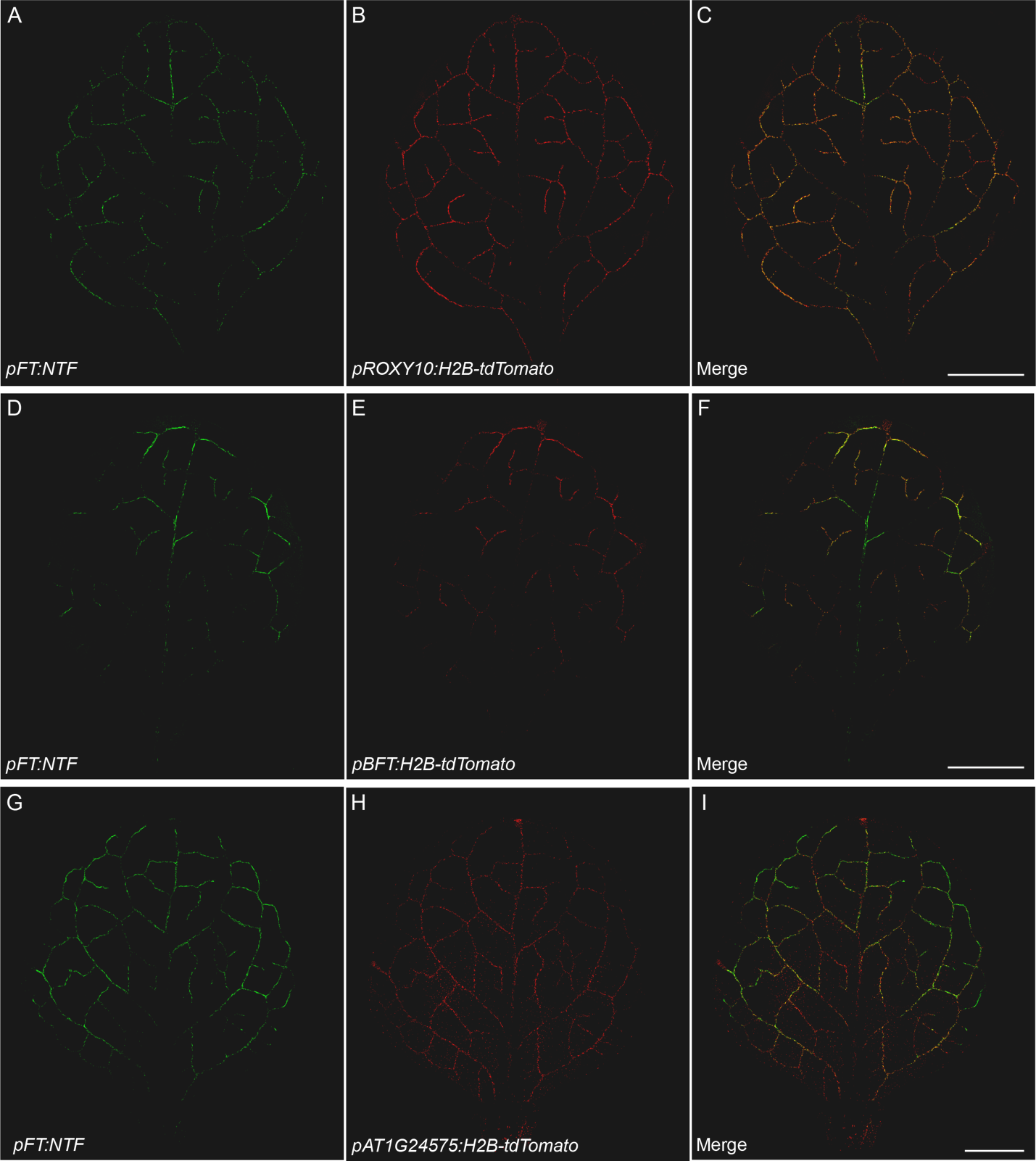

**Supplemental Figure S12.** Spatial expression patterns of GFP signal derived from the *pFT:NFT* (A, D and G) and H2B-tdTomato signals controlled by the promoters of representatives of cluster 7-enriched genes: *ROXY10* (B), *BFT* (E), and *AT1G24574* (H) in true leaves. Yellow color shows an overlap between green and red signals (C, F and I). Scale bar, 1 mm.

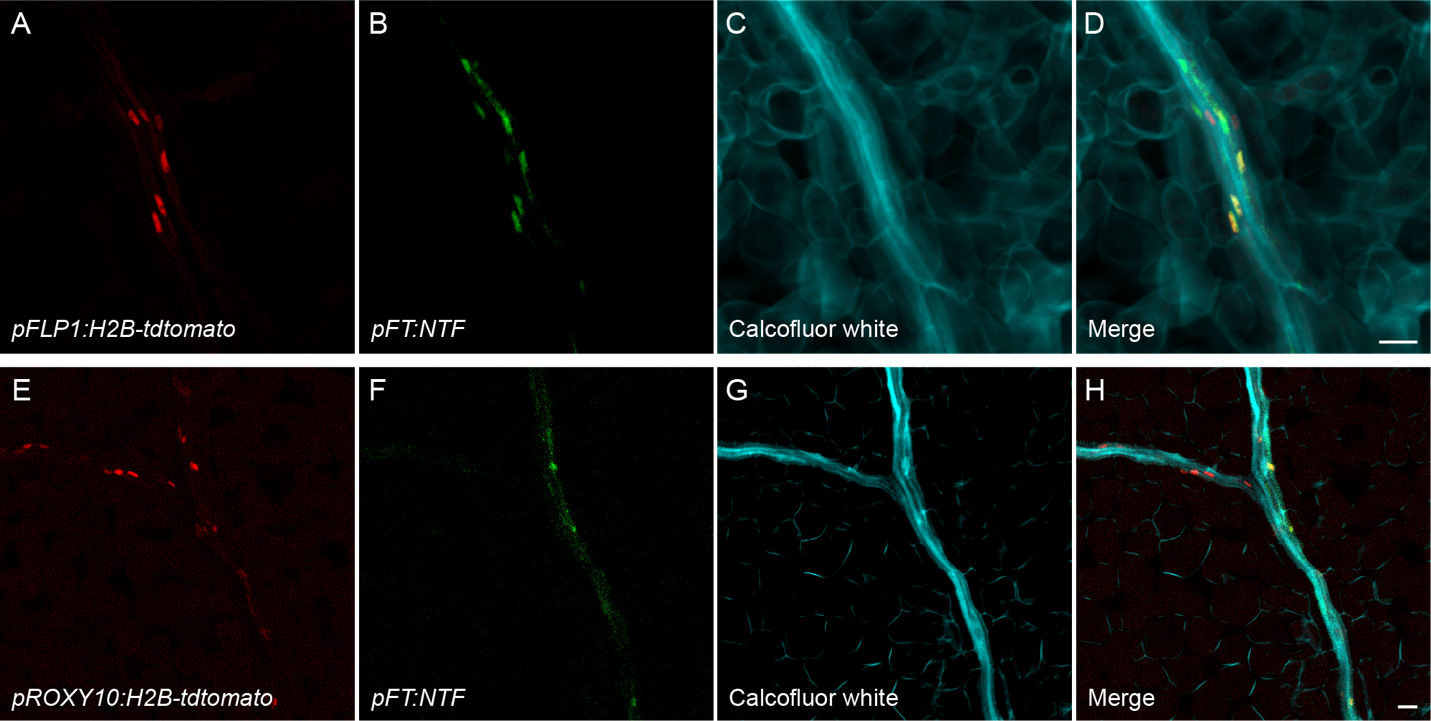

**Supplemental Figure S13.** Enlarged images of phloem companion cells showing spatial expression overlap between *FT* and cluster 7 genes. Images of minor leaf veins of *pFLP1:H2B-tdTomato/pFT:NTF* (A–D), and *pROXY10:H2B-tdTomato/pFT:NTF* (E–H). H2B-tomato driven by the promoter of cluster 7 gene *FLP1* (A) and *ROXY10* (E), NTF (B and F), calcofluor white (C and G), and merged images (D and H).

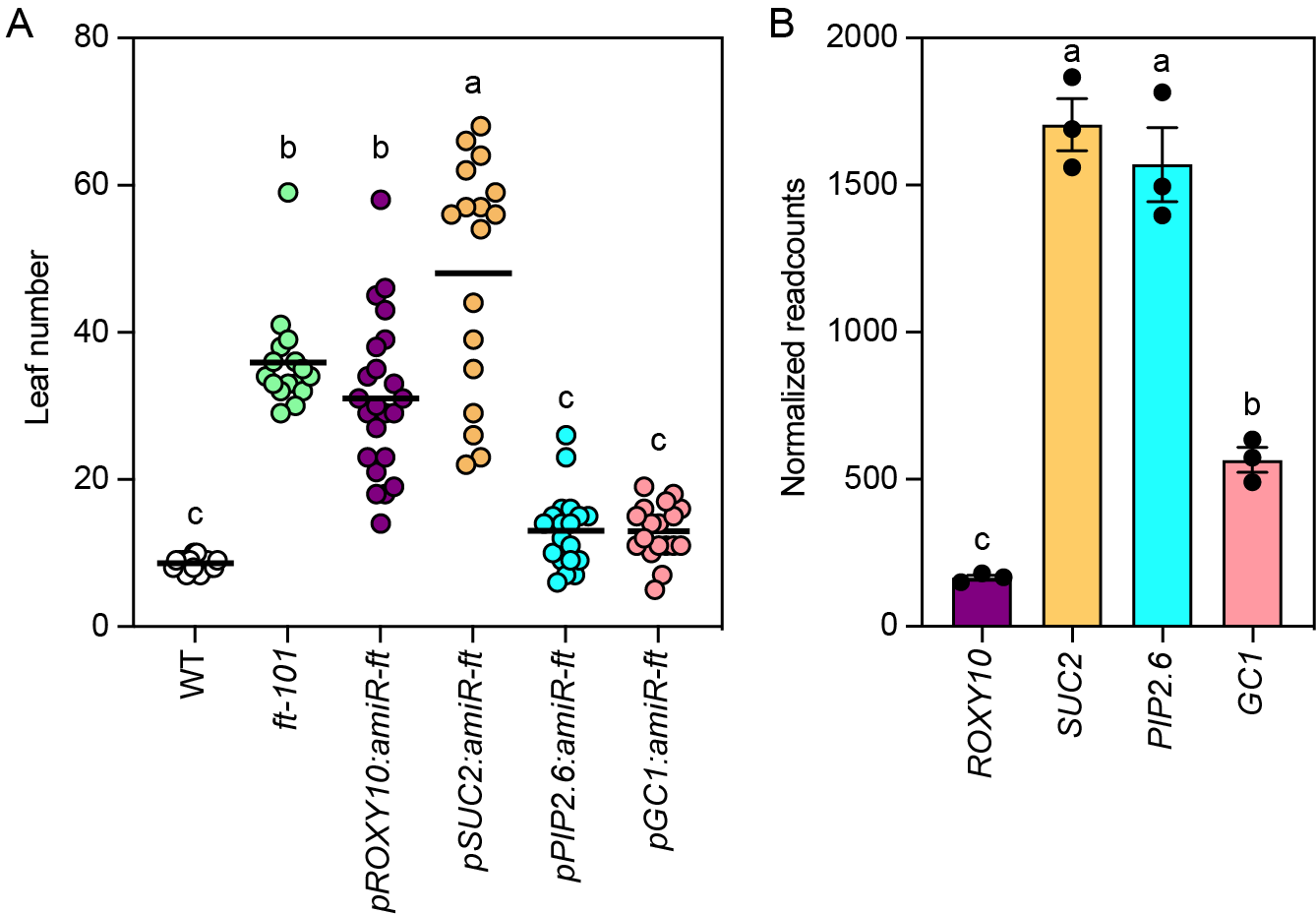

**Supplemental Figure S14.** The silencing of *FT* using amiRNA to confirm the overlap of spatial expression between *FT* and cluster 7-specifc *ROXY10* genes. (A) The effect of amiRNA targeting *FT* mRNA driven by tissue-specific promoters on flowering time. T_1_ plants were grown on hygromycin selection plates for 14 days and transferred to soils in LD+FR. Each dot indicates the flowering time of an individual non-transformant and T_1_ transformant (*n* ≥ 16). Bars indicate mean values, and different letters indicate a statistically significant difference (*P*<0.05, one-way ANOVA and Tukey’s multiple comparison test). (B) Normalized readcounts of *ROXY10*, *SUC2*, *PIP2.6*, and *GC1* genes in wild-type plants in RNA-seq data (Data S10) The results are means ± SEM with each dot representing biological replicates (*n* = 3), and different letters indicate a statistically significant difference (*P*<0.05, one-way ANOVA and Tukey’s multiple comparison test).

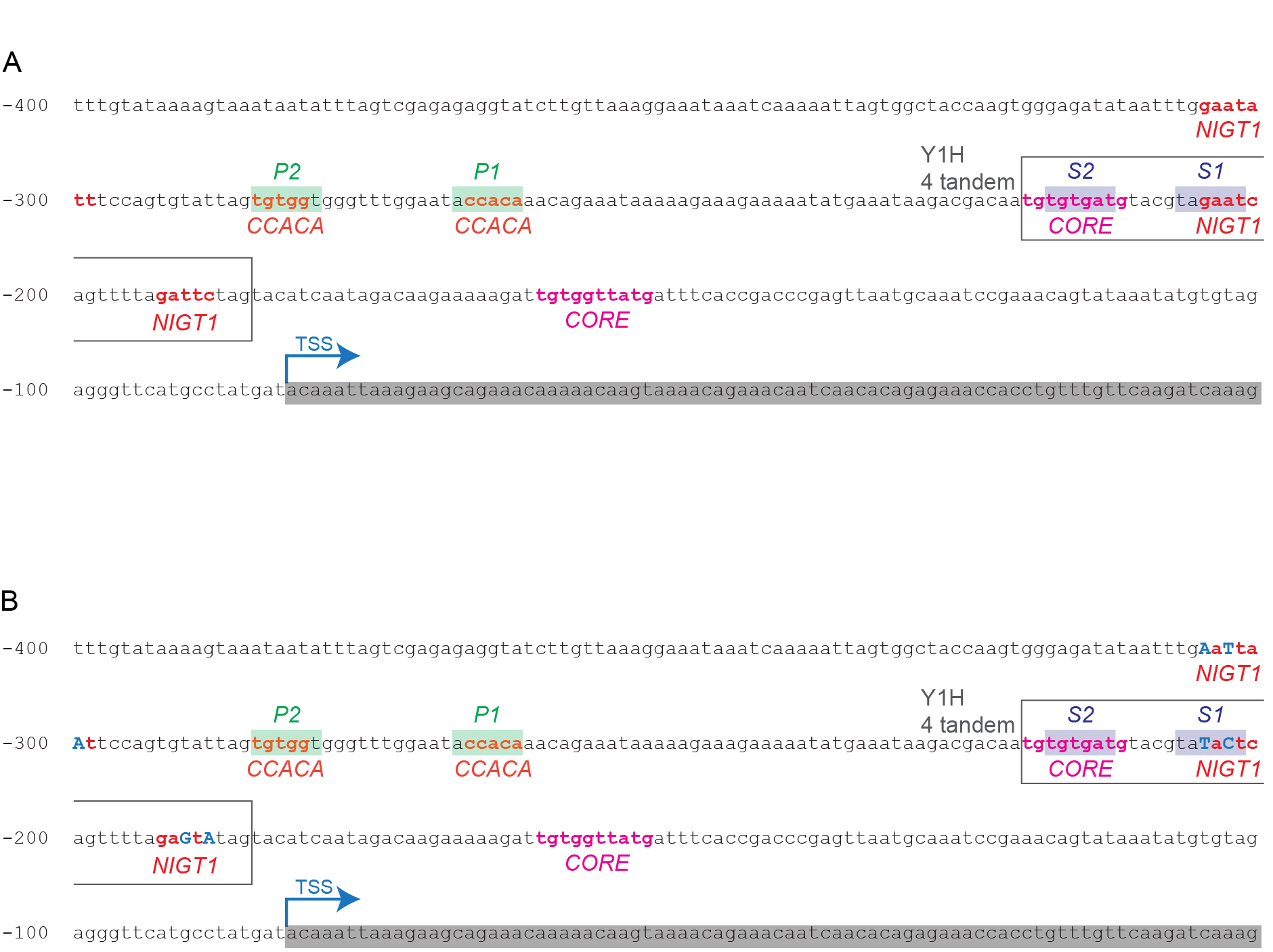

**Supplemental Figure S15.** The 400-bp upstream regions of *FT* promoter sequences (A) and the NIGT-binding site mutated *FT* promoter sequences (B). The position of DNA sequences used for the Y1H screening is marked with a box; four tandem repeats of this sequence were used. The potential NIGT1-binding sites and motifs important for CO-dependent *FT* induction, CO-responsive element (CORE), CCACA, S1/S2, and P1/P2 are indicated. The 5’-UTR is highlighted in gray. The mutations in potential NIGT1-binding sites are indicated with blue capital letters (B).

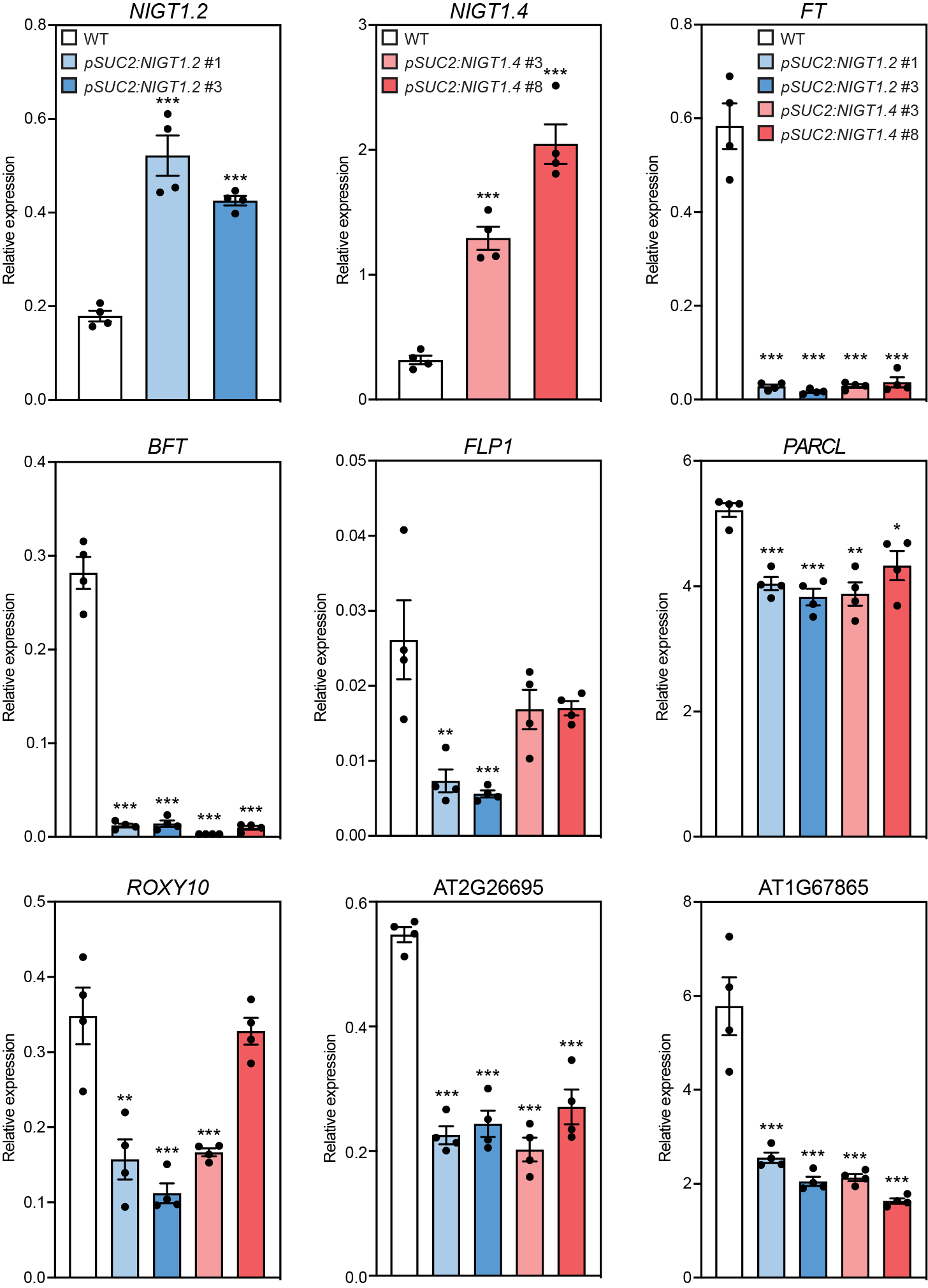

**Supplemental Figure S16.** Relative expression of *NIGT1s* and cluster 7-enriched genes in wild-type (WT), *pSUC2:NIGT1.2*, and *NIGT1.4* lines at ZT4 using quantitative RT-PCR. The results represent the means ± SEM. Each dot indicates a biological replicate (*n* = 4). Asterisks denote significant differences from WT (**P*<0.05; ***P*<0.01; ****P*<0.001, *t*-test).

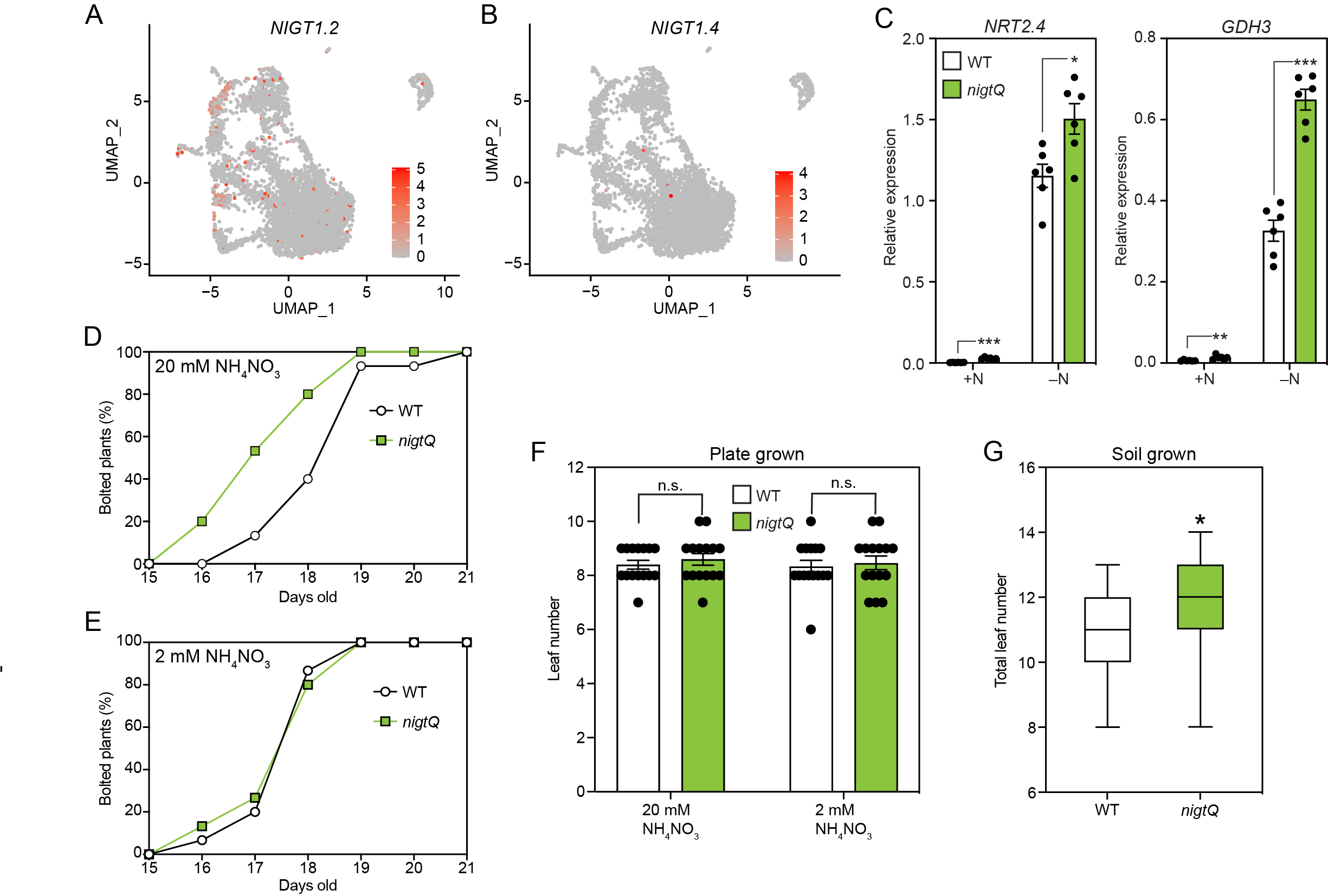

**Supplemental Figure S17.** Expression of *NIGT1.2* and *NIGT1.4* genes and their roles in flowering time. (A and B) UMAP annotated with normalized read counts for *NIGT1.2* (A) and *NIGT1.4* (B). (C) Relative expression levels of nitrogen-deficiency marker genes in 14-day old wild-type (WT) and *nigtQ* seedlings at ZT4. Plants were grown with high nitrogen (+N) and without (–N). (D and E) Percentage of bolted plants grown on 20 mM NH_4_NO_3_ (D) and 2 mM NH_4_NO_3_ containing media (E). Plants were grown on 20 mM NH_4_NO_3_ containing media for 9 days and transplanted to the new media with 20 mM or 2mM NH_4_NO_3_. (F) Leaf numbers of WT and *nigtQ* plants grown on the media with different nitrogen contents at bolting. The results represent the means ± SEM. Each dot indicates a biological replicate (*n* = 15) (n.s., not significant by two-way ANOVA and Tukey’s multiple comparison test). (G) Leaf numbers of WT and *nigtQ* plants grown on nitrogen-rich soil. Asterisks denote significant differences from WT (**P*<0.05, ****P*<0.001, *t*-test).

Table S1 and S2

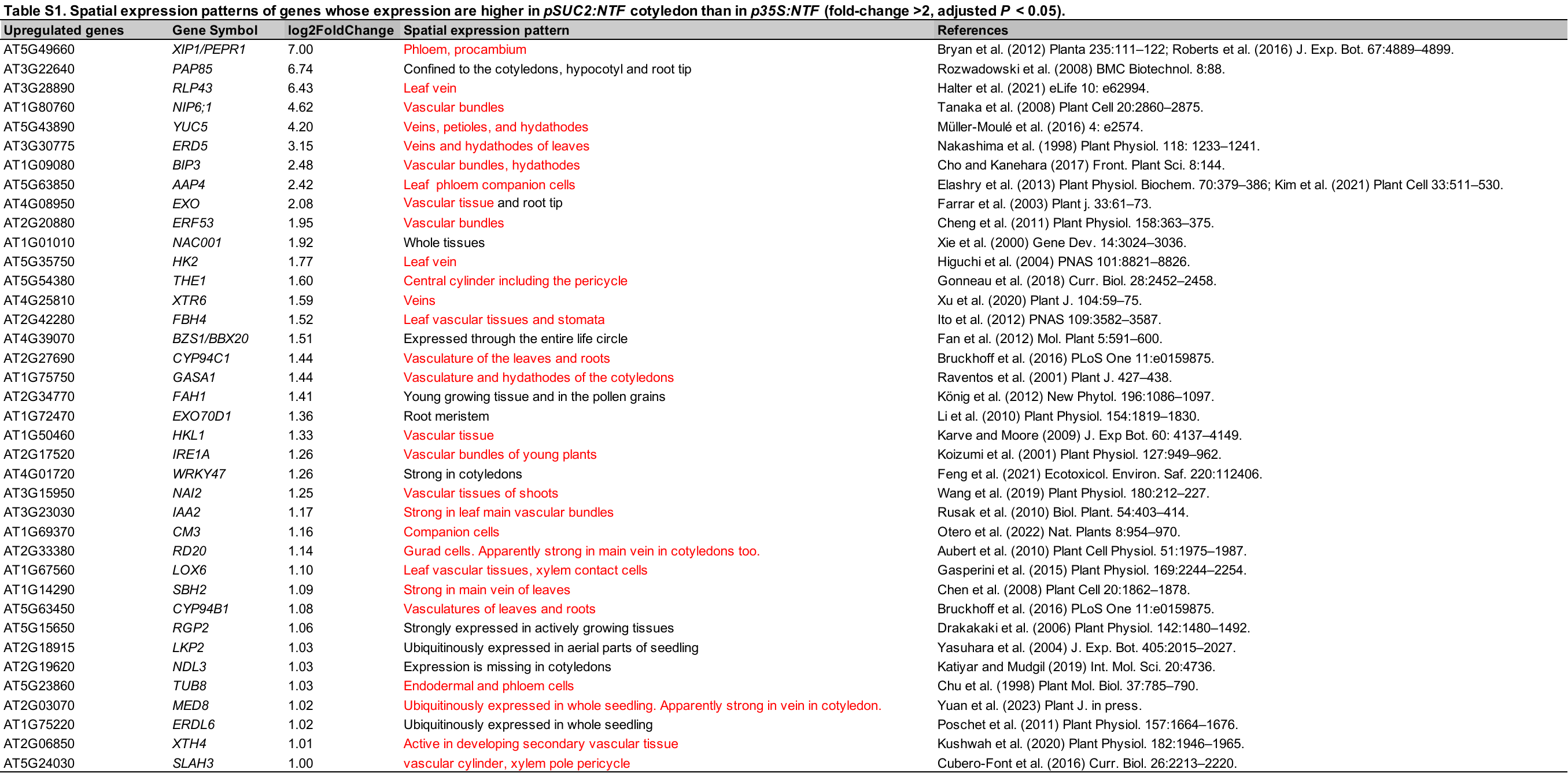

**Table S2.** qRT-PCR primers used in this study.

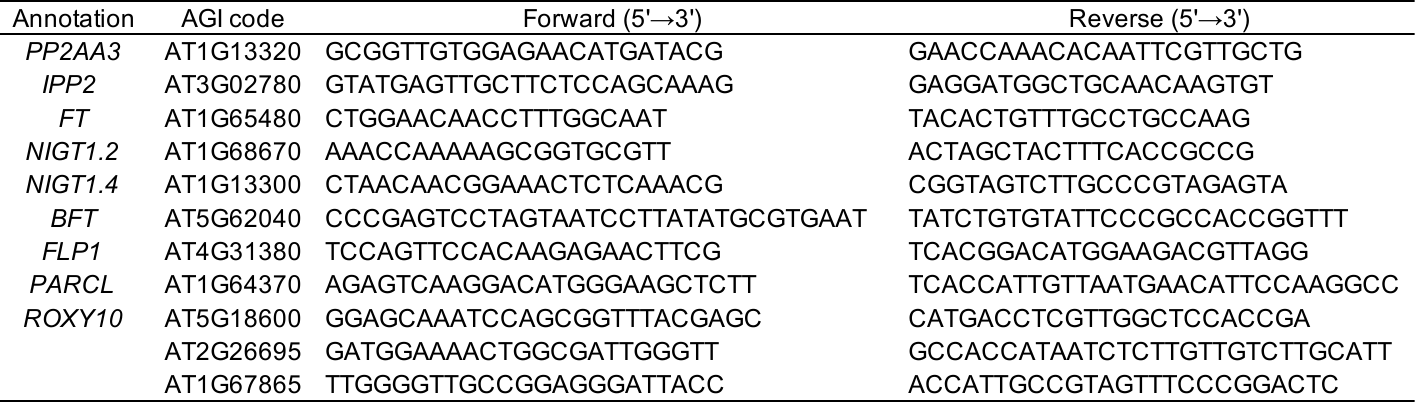
